# pTDP-43 levels correlate with cell type specific molecular alterations in the prefrontal cortex of *C9orf72* ALS/FTD patients

**DOI:** 10.1101/2023.01.12.523820

**Authors:** Hsiao-Lin V. Wang, Jian-Feng Xiang, Chenyang Yuan, Austin M. Veire, Tania F. Gendron, Melissa E. Murray, Malú G. Tansey, Jian Hu, Marla Gearing, Jonathan D. Glass, Peng Jin, Victor G. Corces, Zachary T. McEachin

## Abstract

Repeat expansions in the *C9orf72* gene are the most common genetic cause of amyotrophic lateral sclerosis and familial frontotemporal dementia (ALS/FTD). To identify molecular defects that take place in the dorsolateral frontal cortex of patients with *C9orf72* ALS/FTD, we compared healthy controls with *C9orf72* ALS/FTD donor samples staged based on the levels of cortical phosphorylated TAR DNA binding protein (pTDP-43), a neuropathological hallmark of disease progression. We identified distinct molecular changes in different cell types that take place during FTD development. Loss of neurosurveillance microglia and activation of the complement cascade take place early, when pTDP-43 aggregates are absent or very low, and become more pronounced in late stages, suggesting an initial involvement of microglia in disease progression. Reduction of layer 2-3 cortical projection neurons with high expression of CUX2/LAMP5 also occurs early, and the reduction becomes more pronounced as pTDP-43 accumulates. Several unique features were observed only in samples with high levels of pTDP-43, including global alteration of chromatin accessibility in oligodendrocytes, microglia, and astrocytes; higher ratios of premature oligodendrocytes; increased levels of the noncoding RNA NEAT1 in astrocytes and neurons, and higher amount of phosphorylated ribosomal protein S6. Our findings reveal previously unknown progressive functional changes in major cell types found in the frontal cortex of *C9orf72* ALS/FTD patients that shed light on the mechanisms underlying the pathology of this disease.

## Introduction

Amyotrophic lateral sclerosis (ALS) and frontotemporal dementia (FTD) are progressive neurodegenerative disorders characterized by the loss of neuronal cell populations in the central nervous system (CNS). In ALS, upper motor neurons in the primary motor cortex and lower motor neurons in the spinal cord degenerate, leading to paralysis and respiratory failure typically within 2-5 years of disease onset^1^. FTD is a heterogenous disorder characterized pathologically by the degeneration of the frontal and temporal cortex, leading to progressive cognitive impairments^2^. Despite being symptomatically distinct, ALS and FTD have considerable clinical, genetic, and neuropathological overlap, supporting the notion that these two disorders lie on a disease continuum^3^.

The coexistence of ALS and FTD in the same patients or members of the same family has long been observed and reported in several case studies^4^. Indeed, cross-sectional studies suggest that approximately 50% of ALS patients develop cognitive impairments and ∼30% of patients diagnosed with FTD present with motor neuron symptoms^5^. A multi-center retrospective study found that the order of symptom onset affects survival in ALS-FTD, with ALS onset resulting in shorter survival time. As a major breakthrough in our understanding of ALS and FTD, a G_4_C_2_ hexanucleotide repeat expansion in the gene *C9orf72* was identified as the most common cause of both ALS and FTD^6^. Individuals harboring the repeat expansion can present clinically with ALS, FTD, or both^7^. This variable clinical presentation is also associated with varying disease duration; *C9orf72* patients that present with ALS or ALS-FTD have a median survival of 2.8 and 3 years, respectively, compared to 9 years for patients presenting with FTD only^8^.

A neuropathological hallmark of ALS and FTD is the mislocalization, phosphorylation, and aggregation of TAR DNA binding protein 43 (TDP-43)^9^. TDP-43 is a ubiquitously expressed, nuclear RNA/DNA-binding protein that performs important functions associated with RNA metabolism, including alternative splicing and mRNA stability. The neuropathological confirmation of FTD is referred to as Frontotemporal Lobar Degeneration (FLTD) and positive phosphorylated TDP-43 (pTDP-43) immunoreactivity distinguishes FTLD-TDP from other FTLD pathologies. Specifically, the FTLD-TDP Type B pathology, defined by cytoplasmic pTDP-43 inclusions in neurons of cortical layers II-V and oligodendroglia in white matter^9–11^, is most often observed in *C9orf72* cases that develop clinical features of FTD and ALS. Present in approximately 95% of ALS cases and ∼50% of FTD cases, pTDP-43 burden has been shown to correlate with degeneration of affected cell populations in both ALS and FTD^12–14^. In *C9orf72* carriers specifically, semi-quantitative analyses suggest that the extent of TDP-43 pathology in an affected CNS region correlates with clinical phenotypes^15^. However, the molecular changes associated with quantitative measurements of relative pTDP-43 abundance in a disease-relevant brain region for all frontal cortical cell types have not previously been explored. Given the variability in symptom onset (ALS vs FTD) and discordant timing of clinical progression between ALS and FTD despite a shared genetic etiology, postmortem samples from *C9orf72* ALS/FTD donors with quantitative measurements of pTDP-43 abundance provide a unique opportunity to identify molecular cascades that promote and/or result from TDP-43 dysfunction.

Here we use multiome single-nucleus analysis of postmortem human brain cortex tissue from 26 *C9orf72* ALS/FTD patients and cognitively healthy age-matched controls to gain a more complete picture of the cellular and molecular events altered in different cell types and stages of disease progression based on pTDP-43 abundance. Loss of neurosurveillance microglia and LAMP/CUX2+ cortical projection neurons is observed in donors in early and late stages of disease progression with different levels of pTDP-43. Interestingly, global changes in chromatin accessibility were observed in samples with high levels of pTDP-43, specifically in non-neuronal cells. Donors with high levels of pTDP-43 exhibit several distinct features in non-neuronal cells compared to donors in early stages. The frontotemporal cortex of the patients contains premature oligodendrocytes in which genes encoding for myelin components are downregulated. Other changes include alteration of chromatin accessibility at sites harboring motifs for transcription factors involved in glial cell differentiation, upregulation of ribosomal protein S6 kinase, and increased abundance of phosphorylated ribosomal protein S6. Based on these observations, we propose a sequential cascade of alterations in the regulatory landscape of *C9orf72* ALS/FTD, highlighting the contribution of pTDP-43 accumulation during the progression of neurodegeneration in cell types affected in FTD.

## Results

### Single nucleus multiome analysis of the human dorsolateral prefrontal cortex from *C9orf72* ALS/FTD donors

To investigate molecular changes taking place in specific cell types of the dorsolateral prefrontal cortex of *C9orf72* ALS/FTD patients with different burden of pTDP-43, we utilized a multiome approach to simultaneously analyze the transcriptome and epigenome of Brodmann area 9 of postmortem human brain tissue from *C9orf72* ALS/FTD donors (n=19) and age/sex-matched, healthy controls with normal cognitive and motor function (n=7) (**Fig. 1a** and **Supplementary Table 1**). The average ages are 71 and 69 for control and *C9orf72* ALS/FTD donors, respectively. Fourteen samples were obtained from the Goizueta Emory Alzheimer’s Disease Center Brain Bank and 12 samples were obtained from the Mayo Clinic Brain Bank; we will refer to them as Emory and Mayo cohorts, respectively. All *C9orf72* ALS/FTD donors have a clinical diagnosis of ALS and/or FTD, with neuromuscular abnormalities and different degrees of cognitive impairment. Additional information for each case including age, sex, and co-pathologies is listed in **Supplementary Table 1**. Quantitative measurements of pTDP-43 abundance were performed on all samples using Meso Scale Discovery (MSD) immunoassay in lysates from the same cortical tissue used for multiome analysis. *C9orf72* ALS/FTD donors were then grouped into terciles, referred to as TDPneg, TDPmed, and TDPhigh, based on pTDP-43 levels (**Fig. 1b and Supplementary Table 1**). The presence of cytoplasmic pTDP-43 aggregates in the dorsolateral prefrontal cortex is the defining neuropathological hallmark of FTLD-TDP and it has been reported to associate with more rapid cognitive decline and often found in patients with dementia but not in patients with mild cognitive impairment^16^. Therefore, levels of pTDP-43 were also confirmed using immunohistochemistry for each TDP donor group (**Fig. 1c**).

**Fig. 1.**
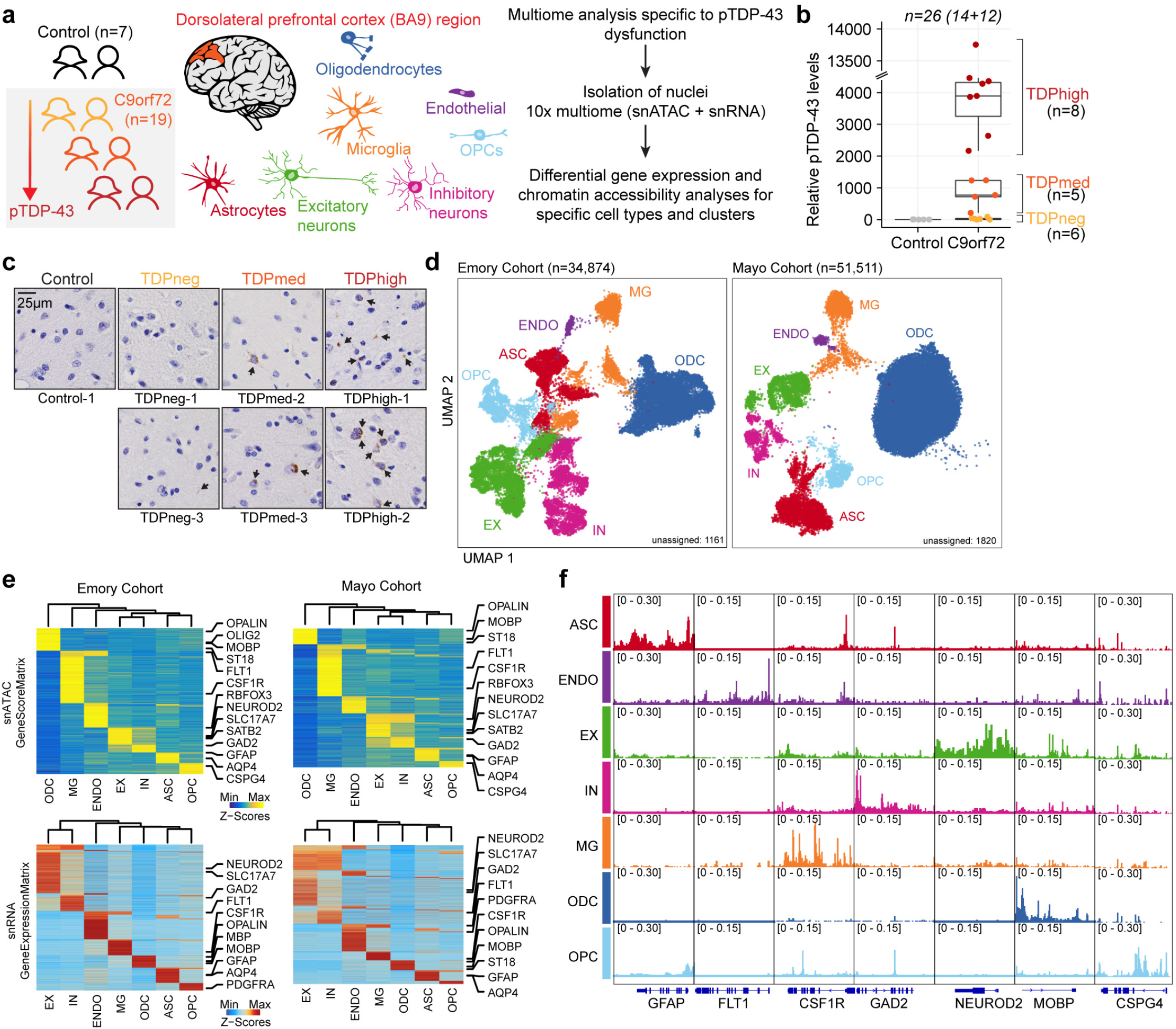
Multiomic single-nucleus analyses identify diverse cortical cell types in the dorsolateral prefrontal cortex of controls and *C9orf72* ALS/FTD patients with different levels of pTDP-43. (a) Schematic representation of single-nucleus multiome profiling (snATAC-seq and snRNA-seq in the same nuclei) of dorsolateral prefrontal cortex samples from 7 control and 19 *c9orf72* ALS/FTD donors analyzed in this study. (b) pTDP-43 levels in control and *C9orf72* ALS/FTD patient cortical tissues. (c) Evaluation of the presence of pTDP-43 aggregates in *C9orf72* ALS/FTD patient cortical tissues and controls. (d) snATAC-seq and snRNA-seq integrated UMAP visualization of major cortical cell types in samples from the Emory (left) and Mayo (right) cohorts, where each dot corresponds to each of the nuclei profiled simultaneously for transcriptome and chromatin accessibility using the 10x multiome platform. (e) Row-normalized single-nucleus gene expression (top) or gene score (bottom) heatmaps of cell-type marker genes for Emory (left) and Mayo (right) cohorts. (f) Pseudo-bulk chromatin accessibility profiles for each cell type at cell-type marker genes in the Emory cohort.

We first analyzed the multiome datasets from all 26 samples; however, we observed a strong batch effect between two groups of samples correlated with the brain bank of origin. Therefore, we separately analyzed the samples from each cohort and performed parallel analyses on each set. We obtained a total of 34,874 and 53,331 single-nucleus multiomes (snRNA-seq and snATAC-seq) from the Emory and Mayo cohorts, respectively, after quality control filtration using the ArchR multiome pipeline^17^ and Seurat snRNA-seq guidelines^18^ (see Methods; **Fig. 1d**; **Supplementary Fig. 1-2**; **Supplementary Table 2**). Dimensionality reduction was performed for each snRNA-seq and snATAC-seq dataset using the ArchR optimized iterative LSI method^17^ and batch effect correction for all samples was performed using Harmony^19^. Uniform manifold approximation and projection (UMAP) and unsupervised clustering with Seurat^20^ were applied to the combined snATAC-seq and snRNA-seq, resulting in a total of 31 distinct cell clusters for Emory samples and 20 distinct cell clusters for Mayo samples, excluding unassigned clusters (**Fig. 1d and Supplementary Fig. 3**). One possible pitfall of droplet based single cell RNA sequencing techniques is the potential inclusion of cell-free RNA, which is commonly referred to as ambient RNA contamination. Using SoupX^21^, we found that there is minimal ambient RNA contamination (**Supplementary Fig. 4a**). Gene activity scores derived from snATAC-seq chromatin accessibility at proximal promoter regions were used to identify marker genes in each cell cluster (**Fig. 1d-f; Supplementary Fig. 3b,d; Supplementary Table 3**). A total of seven major cortical cell types were identified for both cohorts (**Fig. 1d-f; Supplementary Fig. 3**). For the Emory dataset, we performed additional subclustering analysis for each major cell type. Therefore, we identified excitatory neurons (EX; 6576 nuclei and 9 clusters for the Emory cohort; 2570 nuclei and 5 clusters for the Mayo cohort), inhibitory neurons (IN; 4501 nuclei and 7 clusters for the Emory cohort; 1407 nuclei and 5 clusters for the Mayo cohort), astrocytes (ASC; 4,437 nuclei and 6 clusters for the Emory cohort; 4165 nuclei and 3 clusters for the Mayo cohort), microglia (MG; 3255 nuclei and 4 clusters for the Emory cohort; 2580 nuclei and 2 clusters for the Mayo cohort), oligodendrocytes (ODC; 12746 nuclei and 4 clusters for the Emory cohort; 29789 nuclei and 4 clusters for the Mayo cohort), oligodendrocyte progenitor cells (OPC; 2595 nuclei; 3 clusters for the Emory cohort; 1270 nuclei and 2 clusters for the Mayo cohort), and endothelial cells (ENDO; 337 nuclei; 1 cluster for the Emory cohort; 282 nuclei and 1 clusters for the Mayo cohort) (**Supplementary Fig. 3f; Supplementary Table 2b**). The cell type identification was verified by a module score composed of known cell-type specific marker genes (**Supplementary Fig. 3d**). Notably, the Mayo samples have fewer neuronal nuclei (**Supplementary Fig. 3e**), which could explain the strong batch effect observed when attempting to integrate samples from both cohorts. This observation agrees with previous reports indicating difficulties in obtaining high quality single cell gene expression profiles from nuclei isolated from *C9orf72* FTD frontal cortex samples obtained from the Mayo Clinic Brain Bank^22^. Therefore, in the rest of the manuscript, we treat the Emory cohort as the primary dataset and use results obtained with the Mayo cohort to test the validity of the major findings, except for neurons, which are present in low numbers in the Mayo samples.

### Chromatin accessibility was significantly altered specifically in oligodendrocyte lineage cells, microglia and astrocytes from *C9orf72* ALS/FTD donors with high levels of pTDP-43

To address whether the transcriptome and chromatin accessibility are altered progressively and correlate with pTDP-43 levels in each cell type of *C9orf72* ALS/FTD cortex tissue, we first performed systematic pair-wise comparison of gene expression and chromatin accessibility between controls and different groups of *C9orf72* ALS/FTD donors in each of the seven major cell types and cell-type specific clusters observed in the Emory cohort (**Fig. 2**; **Supplementary Table 4**). We first investigated alterations in chromatin accessibility between *C9orf72* ALS/FTD and control samples. A total of 404,124 reproducible 501 bp peaks of chromatin accessible regions were identified in the snATAC-seq dataset. Utilizing the pseudobulk method, we aggregated fragment counts per sample-cell type combination and analyzed them using DESeq2^23^ with multi-factor design to assess the significance of differential regions between control and *C9orf72* ALS/FTD samples with varying levels of pTDP-43. While this method does not adjust for variability among nuclei within the same sample, it offers increased robustness in capturing variations between samples within the same pTDP-43 level group, reducing the likelihood of false positives. A total of 3500 differentially accessible regions (DARs) were identified (**Fig. 2a; Supplementary Table 4a**). Interestingly, the majority of the DARs were observed when comparing control samples with TDPhigh samples (**Fig. 2a**), suggesting that alteration of transcription factor binding might be a hallmark of late disease stages that correlates with increased TDP-43 aggregation and thus reduction in nuclear TDP-43. Surprisingly, DARs are more frequently found in non-neuronal cells, primarily in oligodendrocytes (**Fig. 2b**). The frequent observation of oligodendroglial cytoplasmic pTDP-43 inclusions in *C9orf72* brain tissues with FTLD-TDP^10^ suggests the possibility that changes in chromatin accessibility observed in oligodendrocytes with high levels of cortical pTDP-43 could be due to direct or indirect effects of pTDP-43 accumulation and/or reduction of nuclear TDP-43. Upon analyzing the presence of transcription factor binding motif sequences beneath the summits of these DARs in non-neuronal cells, we observed that motifs associated with transcription factors involved in cell differentiation were prevalent (**Fig. 2c**). Notably, we identified motif sequences for EGR1, KLF5 and ZNF263 in DARs found in all non-neuronal cell types, including microglia, astrocytes, oligodendrocyte precursor cells, and oligodendrocytes (**Fig. 2c**). In contrast, NFIC motif sequences were predominantly present in terminally differentiated glial cells (MG, ASC, ODC), but not in OPCs. Furthermore, CTCF emerged as one of the prominent TF motifs in DARs present in microglia, OPCs, and ODCs. Notably, SOX10, crucial for oligodendrocyte specification, was enriched in DARs identified in oligodendrocyte lineage cells. These findings suggest that the commonly observed abnormalities in glial cells, such as astrogliosis, microglial dysfunction, and oligodendrocyte dysregulation, may arise from widespread changes in transcription factor occupancy in *C9orf72* ALS/FTD. These changes appear to be more prevalent in the presence of pTDP-43 aggregates in the frontal cortical region.

**Fig. 2.**
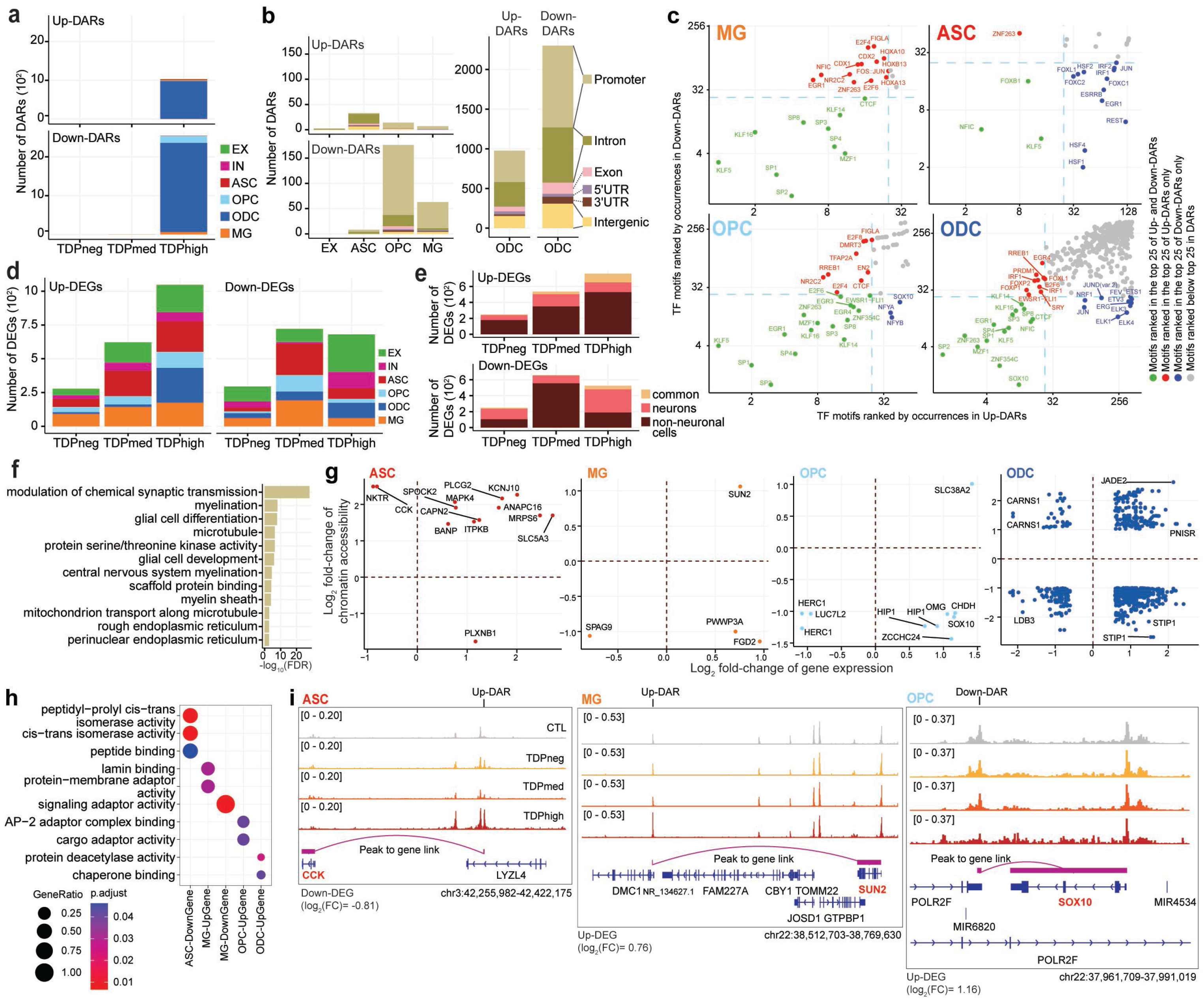
Cell-type specific dysregulation of gene expression and chromatin accessibility in *C9orf72* ALS/FTD donors. (a) Total number of DARs categorized by cell type and pTDP-43 donor group. (b) Total number of DARs found in pTDPhigh donor group categorized by cell types by cell type (left) and distribution of DARs related to functional annotations in oligodendrocyte lineage cells (right). (c) Occurrence of Transcription Factor (TF) motifs in DARs in non-neuronal cells found in the pTDPhigh donor group ranked by the frequency of occurrence. The TF with motif occurrences ranked highest in both Up- and Down-DAR are highlighted in green in the bottom left quadrant of each panel. TFs with motif occurrences ranked highest only in Up-DARs are marked in red and displayed in the top left quadrant, while TFs with motif occurrences ranked highest only in Down-DARs are marked in blue and displayed in the bottom right quadrant (d) Number of DEGs categorized by cell type and pTDP-43 donor group. (e) Number of differentially expressed genes common among neuronal and non-neuronal cells for each pTDP-43 donor group. (f) Gene Ontology enrichment analysis of genes located near DARs found in the pTDPhigh donor group. (g) DEGs with a linked DAR in astrocytes, microglia and oligodendrocyte lineage cells. (h) Gene Ontology enrichment analysis of DEGs with linked DARs as shown in panel (g). (i) Genome track visualization of the *CCK* (chr3:42,255,982-42,422,175), *SUN2* (chr22:38,512,703-38,769,630) and *SOX10* (chr22:37,961,709-37,991,019) loci in astrocytes, microglia and oligodendrocyte precursor cells, respectively.

We proceeded to perform rigorous comparison of transcriptomes to identify alterations between control and *C9orf72* samples. Initially, when analyzing the snRNA-seq and snATAC-seq datasets separately using UMAP, a distinctive batch effect surfaced in the snRNA-seq data, whereas the paired snATAC-seq data remained unaffected (**Supplementary Fig. 2a,b, top panels**). To mitigate this issue, we applied Harmony^19^ batch correction, adjusting for sample, groups categorized by pTDP-43 levels, and sample preparation batch covariates, effectively eliminating the batch effect in both individual datasets and the combined datasets (**Supplementary Fig. 2a,b,c, bottom panels**). For robustness and to reduce false positives in gene expression analysis, we employed the linear mixed-effect model implemented in MAST^24^. This allowed us to meticulously consider technical covariates using the generalized mixed-effect models and model cells individually using the two-hurdle model implemented in MAST (see Methods). Differentially expressed genes were identified across all *C9orf72* ALS/FTD donor groups and cortical cell types, with the greatest number observed in the TDPhigh donor group (**Fig. 2d**; **Supplementary Table 4b).** These findings suggest that both the transcriptome and epigenome are most affected during the late disease stages characterized by high levels of pTDP-43. Interestingly, numerous differentially expressed genes are altered in both neurons and non-neuronal cells. However, a higher number of genes show differential expression in non-neuronal cells (**Fig. 2d-e**), consistent with the findings from the differentially accessible region analysis. These differentially expressed genes are involved in synaptic transmission, myelination, and encode for microtubule proteins. (**Fig. 2f**; **Supplementary Table 4c**).

To leverage the paired multi-omics datasets, we analyzed genome-wide peak-to-gene links utilizing the integrated single-nucleus RNA-seq and ATAC-seq data captured simultaneously in our study. The strength and specificity of each link (as depicted in **Fig. 2i**) are determined by the correlation between chromatin accessibility levels at a given peak and gene expression levels for a specific gene in a single cell. Therefore, a peak and a gene are considered linked if both are altered in the same cell. This approach enabled us to assess how DARs, presumably present at transcriptional regulatory elements, might influence the differential expression of genes in a cell type-specific manner. We found a significant number of DARs linked to DEGs (**Fig. 2g**). Specifically, we found enrichment of differentially expressed genes with linked DARs involved in peptide binding, lamin binding, and chaperone binding in astrocytes, microglia, and oligodendrocyte lineage cells (**Fig. 2h**). For instance, the downregulation of the *CCK* gene is associated with an up-regulated DAR in TDPhigh samples in astrocytes, the upregulation of *SUN2* is associated with an up-regulated DAR in TDPhigh samples in microglia, and the upregulation of *SOX10* is associated with a downregulated DAR in TDPhigh samples in oligodendrocytes (**Fig. 2i**). This analysis suggests that DARs may act as both transcriptional enhancers and silencers in different genomic contexts and cell types.

### *C9orf72* ALS/FTD is associated with impaired oligodendrocyte maturation in late disease stages

Seven distinct cell populations were identified in the oligodendrocyte lineage in the Emory cohort (n=15,341 nuclei), encompassing the largest cell population in the multiome dataset, including oligodendrocyte precursor cells (OPCs) and differentiated oligodendrocytes (ODCs) (**Fig. 3a,b** and **Supplementary Fig. 3**). Oligodendrocyte lineage cells are also the largest cell population in the Mayo cohort (**Fig. 1d**), where six distinct cell populations were identified (**Fig. 3f**). Oligodendrocytes function in the central nervous system by establishing the myelin layer and providing metabolic support to neurons. Importantly, grey matter demyelination has been observed in the motor cortex and the spinal cord of ALS patients^25^. Furthermore, pTDP-43 inclusions in oligodendrocytes are a characteristic neuropathological finding in brains of *C9orf72* ALS/FTD patients^9,10^. These observations suggest that oligodendrocyte dysfunction plays an important role in *C9orf72* ALS/FTD. We first examined the oligodendrocyte lineage clusters from the Emory cohort. Clusters OPC-1, OPC-2, and OPC-3 contain oligodendrocyte precursor cells (OPCs) with high expression of *PDGFRA* and *CSPG4* (**Fig. 3d,e**). The remaining four oligodendrocyte clusters are differentiated oligodendrocytes with higher levels of *OPALIN* and *PLP1*. We noticed that there are proportionally more cells in ODC clusters ODC-C2 and ODC-C3 in TDPhigh compared to other donor groups and controls (**Fig. 3b**). Specifically, an average of 25% of oligodendrocyte lineage cells in TDPhigh donors are found in the ODC-2 cluster (**Fig. 3c**), and less than an average of 2% of oligodendrocyte lineage cells are present in this cluster in TDPmed and TDPneg donors. This suggests that ODC-2 is unique to the late disease stages with high pTDP-43 burden. Based on gene score and hierarchical clustering of marker genes, ODC-C2 cells are transcriptionally distinct from other ODC clusters (**Fig. 3d**), with lower expression of *MOG*, *MOBP*, and *MBP*, which encode for the major protein components of myelin (**Fig. 3e**). ODC-2 cells also have higher expression levels of *TCF7L2* and *ITPR2*, and lower expression of *CNP* and *KLK6* (**Fig. 3e**). Expression of these two genes in ODCs is an indication of newly differentiated premyelinating oligodendrocytes, which is typically a transient stage during adult oligodendrogenesis that survives two days in adult mouse brain^26,27^. The majority of these cells undergo apoptosis while some survive and mature into myelinating oligodendrocytes^28,29^. Therefore, these data suggest that ODC-2 cells represent newly formed premyelinating oligodendrocytes that should not typically be present in high ratios in adult brain, suggesting that either they failed to enter programmed cell death or to proceed into maturation. In contrast to ODC-2, the rest of ODC clusters are composed of mature myelinating ODCs with strong expression of genes involved in myelinating processes^26,30^ (**Fig. 3e**). ODC-3 is also present in large proportion in pTDP-43 high samples (**Fig. 3b**); although cells in this cluster express high levels of myelination genes, their levels are slightly lower compared to ODC-1 and ODC-4. These normal mature myelinating ODCs are found mainly in TDPmed and TDPneg donor groups in earlier disease stages (**Fig. 3b**). Interestingly, compared with oligodendrocytes from control donors, oligodendrocytes from the TDPhigh donor group exhibited downregulation of MOG, the myelin oligodendrocyte glycoprotein (**Fig. 3f** and **Supplementary Table 4**). When we independently analyzed oligodendrocyte lineage clusters from the Mayo cohort (**Fig. 3g**), the Mayo ODC-1 cluster exhibits the same premature premyelinating oligodendrocyte markers, with high expression of *TCF7L2* and *ITPR2* and lower expression of *CNP* and *KLK6* compared to other clusters (**Fig. 3h**). Similar to the Emory cohort, there are proportionally more cells in the Mayo ODC-1 cluster in TDPhigh compared to controls samples (**Fig. 3i-j**) and oligodendrocytes from the Mayo TDPhigh donor group exhibit downregulation of the same myelin-associated genes (**Fig. 3k and Supplementary Table 4**). These findings, observed in the two cohorts studied, further strengthen the conclusion that a large portion of oligodendrocytes in the dorsolateral prefrontal cortex in late FTD disease stages with high pTDP-43 burden remain in the typically transient premyelinating stage and defective in myelination. We also found down-regulated DARs located near the promoter region of genes involved in myelination (**Fig. 3l and Supplementary Table 4**), including *OPALIN*, *MAG*, *PLLP* and *MOBP*. However, decrease in chromatin accessibility at the promoter regions of these genes does not correlate with statistically significant changes in steady-state RNA levels based on the linear mix-effect model. mRNAs encoded by these genes are bound by TDP-43 in the mouse brain^31,32^ and the development of oligodendrocytes has been shown to be regulated by TDP-43^33^. Combining our new findings with previously published data, we hypothesize that the downregulation of these genes could be a direct consequence of the loss of nuclear TDP-43 in the late disease stage, accompanied by cytoplasmic accumulation of pTDP-43. Interestingly, the expression of these myelin-associated genes was not significantly altered in AD donors with early or late pathology^34^, suggesting myelination impairment might be a unique defect associated specifically with pTDP-43 pathology.

**Fig. 3.**
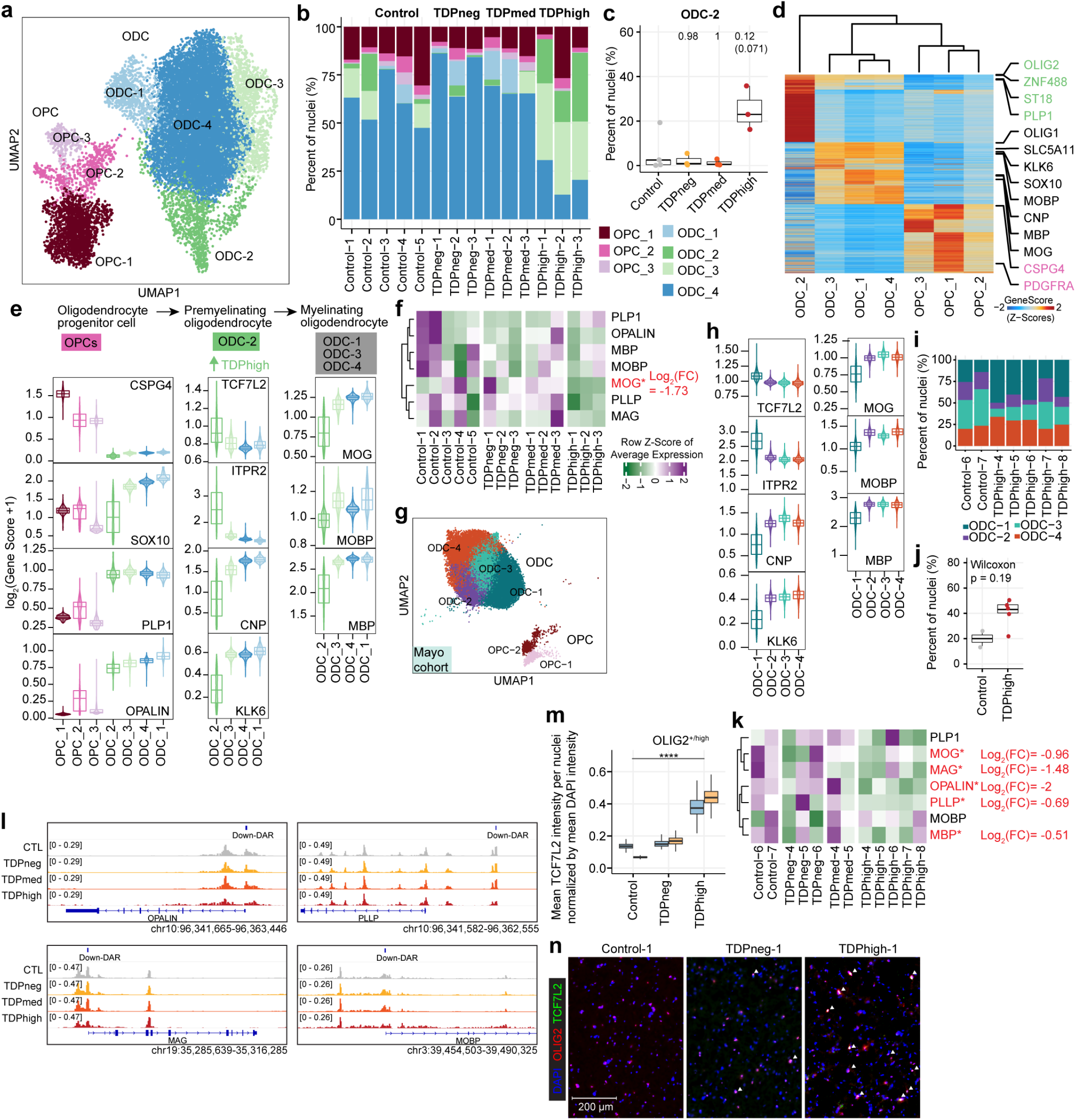
Premature and premyelinating oligodendrocytes are unique to high pTDP-43 donors in late disease stages. (a) UMAP plot of oligodendrocyte lineage cells for Emory samples. (b) Proportion of oligodendrocyte precursor cells (OPC) and oligodendrocytes (ODC) clusters in each sample, including donors with different levels of pTDP-43 and cognitively normal controls of the Emory cohort. (c) Proportion of ODC-C6 in different Emory cohort pTDP-43 donor groups (Kruskal–Wallis test with Benjamini–Hochberg correction (p-adj=0.0771) and without correction (p-value=0.0129). (d) Plot of snATAC-seq gene scores ordered by hierarchical clustering with marker genes distinguishing each ODC cell cluster for Emory cohort samples. (e) Illustration of developmental stages of oligodendrocyte lineage cells for Emory samples. Developmental stage specific genes and their gene scores are shown for each cluster (bottom), highlighting the unique characteristics of ODC-C6 with high expression of premyelinating oligodendrocyte genes. (f) Average expression of myelin associated genes in Emory samples. (g) UMAP plot of oligodendrocyte lineage cells for Mayo cohort samples. (h) ODC-1 in Mayo samples exhibit high expression of premyelinating oligodendrocyte genes, similar to ODC-2 in Emory samples. (i) Proportion of ODC-1 in Mayo control and TDPhigh samples (p-value=0.19, Wilcoxon Rank Sum). (j) Proportion of ODC clusters in Mayo cohort control and TDPhigh samples. (k) Average expression of myelin associated genes in Mayo samples. (l) IGV track view of changes in chromatin accessibility in close proximity to promoter regions of myelin associated genes. (m) Quantification mean TCF7L2 intensity per OLIG2-high nuclei normalized by mean DAPI intensity after nuclei segmentation. One-way ANOVA, ****p value < 0.0001.(n) Immunostaining of human postmortem cortical tissue for the oligodendrocyte lineage marker OLIG2 (red), premature oligodendrocyte marker TCF7L2 (green) and DAPI in blue. Overlapping OLIG2 and TCF7L2 staining were marked with white arrowhead.

To validate the high abundance of premature oligodendrocytes found in the TDPhigh samples based on results from the single-cell analysis, we performed immunofluorescence microscopy for TCF7L2, OLIG2, and NeuN, along with proteinase K treatment and photobleaching to remove lipofuscin autofluorescence using control, TDPneg and TDPhigh samples from the Emory cohort. Sections were stained with oligonucleotide-tethered secondary antibodies, followed by gel embedding before imaging. Nuclei were segmented and classified based on OLIG2 and NeuN staining intensities into “OLIG2-high”, “NeuN-high”, or “other” categories (**Supplementary Fig. 5a-c**; see Methods). OLIG2-high nuclei represent cells with oligodendrocyte lineages. Analysis of DAPI-normalized TCF7L2 intensity in OLIG2-high nuclei revealed significantly higher TCF7L2 signals in TDPhigh samples compared to control and pTDPneg samples (**Fig. 3m**). Examination of images from upper cortical regions confirmed the quantification results, showing a higher number of overlapped TCF7L2-positive and OLIG2-positive nuclei (**Fig. 3n**). To ensure the observed results were not artifacts of the proteinase K, photo clearing and oligonucleotide-tethered secondary antibody procedures, we employed a second approach using TrueBlack Lipofuscin Autofluorescence Quencher on paraformaldehyde-fixed floating tissue sections, followed by confocal imaging. Consistently, we found a higher number of OLIG2+/TCF7L2+ nuclei in TDPhigh samples compared to control and TDPneg samples (**Supplementary Fig. 5d**). These independent validation methods confirm an overabundance of OLIG2+/TCF7L2+ nuclei in the grey matter region of pTDPhigh samples, reinforcing the single-cell analysis findings. The discovery of highly abundant premature oligodendrocytes specifically in pTDPhigh samples presents a novel and unique insight into *C9orf72* ALS/FTD disease progression and TDP-43 proteinopathy, not previously reported. Given prior evidence of pTDP-43 inclusions in oligodendrocytes and that mRNAs encoding myelin proteins are bound by nuclear TDP-43^31,32^, we speculate that the presence of pTDP-43 aggregates, or the correlated absence of nuclear TDP-43, may play a direct role in dysregulating mRNAs encoding myelin components, thus affecting the maturation of oligodendrocytes and their ability to myelinate neurons, a possible unique feature to *C9orf72* ALS/FTD patients with significant pTDP-43 burden.

### Loss of frontal cortical neurosurveillance microglia is a hallmark of both early and late stages of *C9orf72* ALS/FTD, and is similarly prevalent in early and late AD

Microglia typically account for 5% of all brain cells^35^ and have the highest expression of *C9orf72* compared to other cortical cell types^36^. In agreement with these observations, we also find that the gene score activity and gene expression of *C9orf72* are highest in microglia compared to other cortical cell types, in samples from both the Emory and Mayo cohorts (**Supplementary Fig. 3f**). As the resident immune cells, microglia are thought to contribute to the increased inflammation reported in the ALS/FTD disease spectrum^37,38^. snRNA-seq studies of human cortical tissues have reported that microglia form one large diffuse cluster, suggesting that, instead of distinct cell types, human microglia populations vary gradually in their transcriptome states^39,40^. A recent ultra deep analysis of human microglia further identified 12 transcriptional states in this cell population; however, these transcriptional states were not captured by independent snATAC-seq performed in the same cohort^41^. We first analyzed microglia from the Emory cohort; 4 cortex cell clusters with a total of 3,438 nuclei have microglia identity and express known microglia markers, including *AIF1*, *RUNX1*, *PTPRC* (CD45), *CX3CR1*, *P2RY12*, *TMEM119*, and *ITGAM* (CD11b)^39^ (**Fig. 4a,b and Supplementary Fig. 3d**). Each of these four microglia clusters exhibits a distinct set of expressed genes, snATAC-seq peaks, and transcription factor binding motifs (**Fig. 4b,c**), suggesting they correspond to distinct identities of microglia cells. It is possible that the ability to discover these four distinct microglia populations is based on using paired snRNA-seq and snATAC-seq from the same nuclei and unsupervised clustering with combined snATAC-seq and snRNA-seq, instead of snRNA-seq or snATAC-seq independently, thus achieving a more defined cell cluster identity. MG-1 is the largest microglia cluster (n=1,751 nuclei) and appears to be in a combination of homeostatic and activating states based on the presence of marker genes in the multiome data. This cluster exhibits the highest expression of microglia homeostatic marker genes^7^, including *CX3CR1*, *TMEM119* and *CSF1R* (**Fig. 4b**). Cells in this cluster also express genes characteristic of the activating state, including inflammatory genes involved in antigen presentation (*CD86*, *CD80*; MHC II – *C1QA*, *C1QB*, *C1QC*), reactive chemokines (*CCL2*, *CCL3*), and interleukin (*IL-1a*, *IL18*) (**Fig. 4b**). TF binding motifs for SPI1 (also known as PU.1), a TF that is essential for microglia activation^42,43^, are specifically enriched in the MG-1 cluster at chromatin accessible regions (**Fig. 4c**). IRF8, another critical TF that transforms microglia into a reactive phenotype^42^, is uniquely highly expressed in the MG-1 cluster. Distinct from MG-1, the other three MG clusters exhibit lower expression of genes involved in microglia homeostasis, suggesting that the shift away from the homeostatic state may be due to downregulation of these genes (**Fig. 4a-c**). Clusters MG-2 and MG-3 exhibit moderate expression of markers for an alternative M2-like microglia state (**Fig. 4b**), which has been proposed to be an anti-inflammatory state that plays a protective role in the brain in contrast to reactive microglia^44^. Specifically, MG-2 and MG-3 are defined by different sets of neurotrophic factors (MG-2: BDNF/GDNF/ NTS; MG-3: BDNF/GDNF/ NGF) (**Fig. 4b**) that have established roles in supporting neuron survival and modulate the formation of long-term memories^45,46^. In addition, MG-2 is marked by genes involved in cell-adhesion, pro-proliferation (*UBE4B*), interferon type I interferon receptor binding, and the Complement receptor gene *CR1* (**Fig. 4b**). In contrast, MG-3 cells are marked by genes encoding serotonin receptors and genes involved in G-protein-coupled receptor signaling. MG-4 is a distinct cluster that expresses markers of microglia and astrocytes, such as *GFAP*, *VCAN,* and *AQP4* (**Fig. 4b** and **Supplementary Fig. 3c**), suggesting this cluster might correspond to a specific subset of microglia cells that is phenotypically transitioning into astrocyte-like cells. This type of cell has been shown to be present in an inherited model of ALS^47^. The expression of *PAX6* in MG-4 further confirms the similarity with glial cells in this cluster (**Fig. 4b**). Samples from the Mayo cohort captured 2723 nuclei with microglia identity forming 2 unsupervised clusters (**Supplementary Fig. 6a**), probably due to fewer microglia captured. However, the distinct groups of microglia cells found in the Emory cohort can also be identified in the Mayo cohort, sharing the same marker genes based on module gene scores (**Supplementary Fig. 6b**). For example, the Mayo MG-1 cluster is the largest microglia cluster (n=2,027 nuclei) and expresses the same marker genes as the Emory MG-1 cluster (**Supplementary Fig. 6b**). Although markers of the other three Emory microglia clusters can be found in different groups of nuclei within the Mayo MG-2 cluster (n=553 nuclei), the separation between these microglia cell types is not as distinctive as in the Emory cohort (**Supplementary Fig. 6b**).

**Fig. 4.**
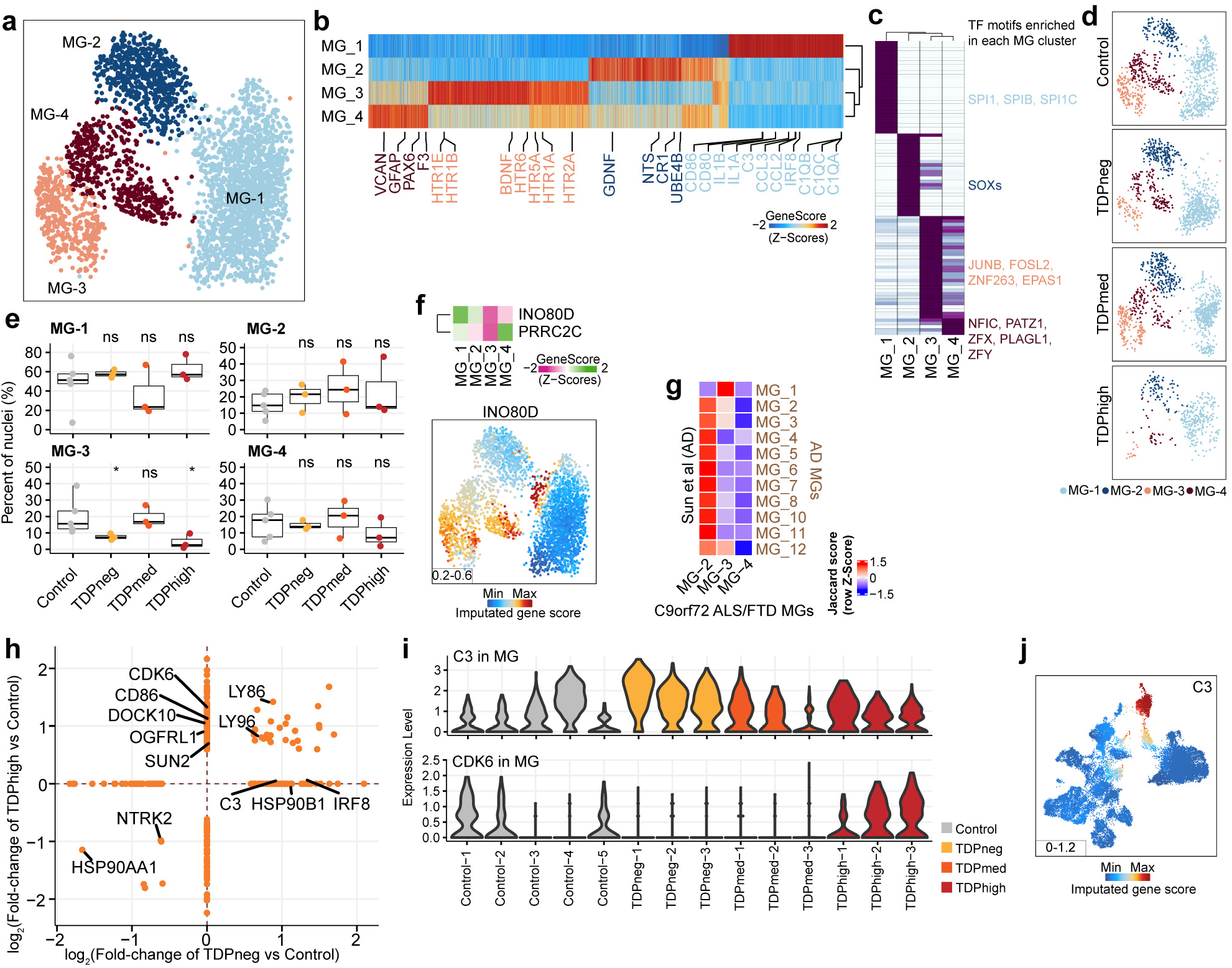
Loss of neuronal surveillance microglia in *C9orf72* ALS/FTD donors with low and high pTDP-43. (a) UMAP plots of the four microglia clusters. (b) Heatmap showing the row-normalized pseudo-bulk gene score in each snATAC-seq cluster split by nuclei from each of the four MG clusters; rows are organized based on hierarchical clustering and the key genes that define the microglia lineages are marked (bottom). (c) Heatmap of motif enrichment at differential marker peaks of each microglia cluster. Color indicates the motif enrichment (−log_10_(P value)) based on the hypergeometric test. TFs specifically enriched for each MG cluster are highlighted using the same cell cluster specific colors as in (a). (d) UMAP plots of the distribution of each pTDP-43 sample group for the four microglia clusters. (e) Fraction of each MG cluster in control and pTDP-43 donor groups (Kruskal–Wallis test with Benjamini–Hochberg correction; p>0.05, n.s.). (f) Gene activity of MG-1 marker genes from the AD dataset. (g) Heatmap showing the similarities of marker genes between Sun et al. and MG-2, MG-3, and MG-4 microglia clusters. The jaccard score indicates the percentage of pairwise overlapping genes. (h) DEGs found in pTDPhigh compared to control and pTDPneg compared to control samples. (i) Violin plots showing gene expression levels of the *C3* and *CDK6* genes in microglia of all Emory samples. (j) Gene activity score for the *C3* gene.

We observed a reduction in microglia in TDPhigh samples from *C9orf72* ALS/FTD donors compared to control samples (**Fig. 4d**). Specifically, the proportion of cells in the MG-3 cluster is significantly lower in both the TDPneg and TDPhigh samples compared to control samples (**Fig. 4e**). However, the proportions of each MG cluster vary in TDPmed samples, likely due samples in the TDPmed group represent a mixture state in pTDP-43 levels. Additionally, there were fewer MG-3 cells in the TDPhigh group compared to the TDPneg group, suggesting that the loss of cells in this cluster is independent of pTDP-43 but could be exacerbated by the high accumulation of pTDP-43 aggregates. However, the changes in TDPmed samples were not significant. Notably, our analysis revealed that the MG-3 cluster shares similar marker genes, such as *INO80D* and *PRRC2C* (**Fig. 4f**), with the MG-1 cluster found in patients with AD^41^, which has been implicated in neuronal surveillance function^48^. This MG-1 cluster shows decreased proportion of cells in both early and late AD stages (see Sun et al.^41^, Fig. 1C). Furthermore, we found that the MG-3 cluster exhibits high expression levels of various neurotransmitter receptors (**Supplementary Fig. 6c**). While MG-1 shares signatures with all AD microglia clusters due to its mixed state (**Supplementary Fig. 6d**), MG-3 demonstrates the closest transcriptome resemblance to the MG-1 cluster in samples from AD donors (**Fig. 4g**). When examining differential gene expression, we find that there are no DEGs encoding neurotransmitter receptors in the TDPneg group compared to controls, whereas the neurotransmitter receptor genes *OGFRL1* and *DOCK10* are upregulated in the pTDPhigh group (**Fig. 4h**). Notably, these two genes are more highly expressed in the MG-1 homeostatic cluster. These data suggest that the reduction in neuronal surveillance microglia occurs in the early disease stage, preceding the accumulation of pTDP-43, perhaps as a consequence of the high expression of the *C9orf72* gene in microglia, although we did not find changes of *C9orf72* expression in microglia between control and *C9orf72* samples. Furthermore, the decrease in the proportion of cells in this cluster becomes more pronounced with high levels of pTDP-43 accumulation, and this trend is also observed in AD donors in late stages with extensive AD pathology.

In addition to the decline in microglia responsible for neuronal surveillance, we observed upregulation of the neuroinflammation genes *C3* and *IRF8* in the pTDPneg group in the pseudobulk microglia analysis as well as in the MG-1 cluster in the Emory samples (**Fig. 4h,i and Supplementary Table 4**). We found similar findings in the Mayo cohort, where *IRF8* is also significantly upregulated in pTDPneg samples (**Supplementary Table 4**). *C3* expression shows a similar trend, although it is not statistically significant (**Supplementary Fig. 6e**). The *C3* gene encodes the complement component 3 protein, crucial in the complement cascade, which plays a significant role in mediating phagocytosis and synapse pruning and elimination in the adult brain^49^. Studies have shown increased levels of C3 protein in early stages of AD, escalating further in advanced stages^50^, suggesting this is another feature shared between early stages of *C9orf72* ALS/FTD and AD. It has been proposed that early synaptic loss in AD is mediated by complement-related pathways and microglia before the accumulation of amyloid plaques. Inhibition of complement component proteins and receptors has been shown to reduce the number of phagocytic microglia and mitigate early synapse loss, thus preserving cognitive function^51^. However, in late-stage AD, the presence of amyloid plaques can trigger the extracellular release of C3 by astrocytes. This released C3 can interact with receptors on microglia and neurons, leading to further synaptic loss, which correlates with more extensive cognitive decline and dysregulation of amyloid plaque phagocytosis^52^. Our results indicate that *C3* is highly expressed in microglia compared to other cortical cell types (**Fig. 4j**) and that upregulation of *C3* is only observed in pTDPneg samples (**Fig. 4i**). This suggests that in the early stages of *C9orf72* ALS/FTD, before the accumulation of pTDP-43, increased *C3* expression and release from microglia could lead to early-stage synapse pruning, similar to what has been proposed in early AD. The activation of *C3* and the complement cascade in early *C9orf72* ALS/FTD could result from repeat expansions in the *C9orf72* gene, which is highly expressed in microglia compared to other cell types. Surprisingly, we did not observe changes in *C3* or *C1QA* in any cell type during the late stage of FTD with extensive pTDP-43 accumulation, suggesting that, although the complement cascade is activated in early and late stages of AD, it may not play a significant role in late-stage *C9orf72* ALS/FTD. Several changes in gene expression in microglia are specific to late disease stages, including upregulation *CDK6*, *CD86*, and *SUN2* in pTDPhigh samples from the Emory cohort (**Fig. 4h,i**). Although *CDK6* was not statistically differentially expressed in the Mayo samples, it shows a similar trend as that observed in the Emory pTDPhigh samples (**Supplementary Fig. 6e**). *CDK6* upregulation has also been observed in lymphoblasts from FTLD patients with mutations in the *GRN* gene^53^, suggesting a shared phenotype between TDP-43 proteinopathy with diverse genetic causes. *SUN2* (also known as UNC84B) and *CD86* are both interferon-stimulated genes activated through Type 1 and Type II IFNγ-stimulated signaling pathways, respectively^54,55^. These observations suggest that, in contrast to early disease stages where the complement cascade is activated in microglia, late disease stage with high levels of pTDP-43 are characterized by the activation of interferon responses.

### Astrocyte dysregulation becomes more pronounced in the advanced stages of the disease characterized by high levels of pTDP-43

Astrocytes (ASCs) represent another cortical cell type known to become reactive and to respond to disease state in neurodegenerative diseases, particularly via dysregulation of metabolic pathways^56^. Cell clusters ASC-1 to ASC-4 with a total of 3,703 nuclei in the Emory cohort and ASC-1 to ASC-3 with a total of 4,162 nuclei in the Mayo cohort can be identified as having astrocyte identity based on high expression of *GFAP*, *AQP4* and *SLC1A2* (**Fig. 5a** and **Supplementary Fig. 3c,d**). Each astrocyte subpopulation exhibits a distinct set of expressed genes (**Supplementary Table 3**) and the ASC-3 cluster in the Emory cohort and the ASC-2 cluster in the Mayo cohort have higher levels of *GFAP*, which encodes the main astrocyte intermediate filament protein and it is a signature of reactive astrocytes^57^, suggesting these two clusters are the most reactive astrocytes in both cohorts. The gene activity of *MT2A*, which encodes a metallothionein protein associated with neuronal injury^58^, is higher in these two reactive astrocyte clusters found in the Emory and Mayo cohorts, and lower in ASC-2 in the Emory cohort (**Fig. 5b**).

**Fig. 5.**
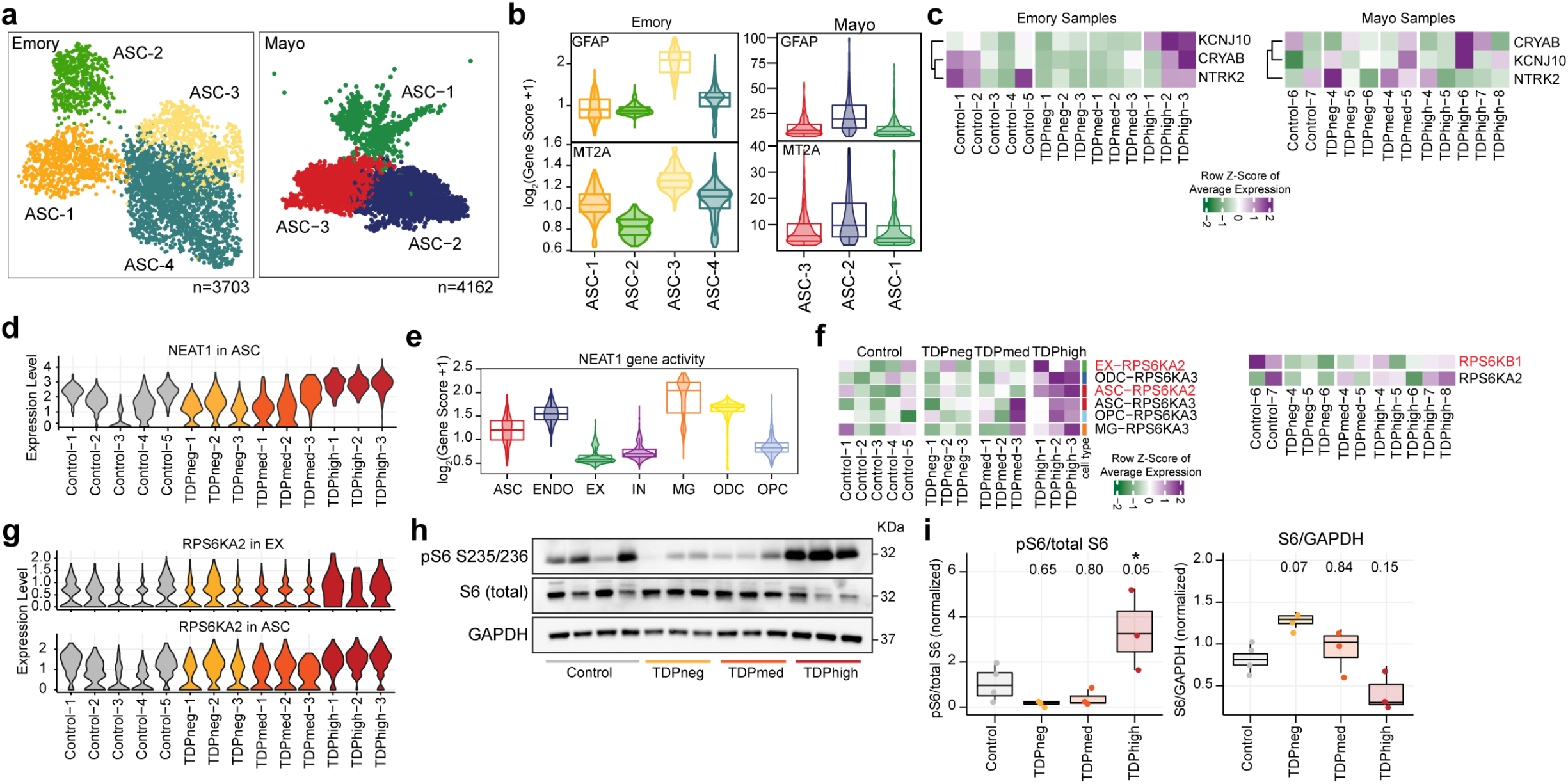
Changes of gene expression in astrocytes are more pronounced in pTDPhigh samples. (a) UMAP plots of astrocyte (ASC) clusters for Emory (left) and Mayo (right) cohorts. (b) Astrocyte clusters exhibit differential levels of *GFAP* and *MT2A* in both cohorts. (c) Changes in gene expression of astrocyte reactivity marker genes. (d) *NEAT1* expression in astrocytes for all samples in the Emory cohort. (e) Cell type specificity of *NEAT1* gene activity. (f) Changes of *RPS6KA2* and *RPS6KA3* gene expression are more significant in pTDPhigh samples. (g) *RPS6KA2* is upregulated both in astrocytes and excitatory neurons. (h) Western immunoblot images of phosphorylated ribosomal protein S6, total ribosomal protein S6, and GAPDH (i) Quantification of the ratio of phosphorylated ribosomal protein S6 to total protein (left) and total ribosomal protein S6 to GAPDH. 1-way ANOVA with Tukey’s post hoc test, adjusted p-values are shown, and *P < 0.05).

We observed various DEGs in astrocytes, distinguishing astrocyte reactivity in early and late disease stages, particularly for samples in the Emory cohort. However, due to the low number of astrocytes in our dataset, we could not perform astrocyte cluster specific differential gene expression analysis using the mixed effect model. Therefore, for the remaining astrocyte analysis, we will discuss changes of gene expression found in all astrocytes using a linear mixed effect model. The astrocyte-reactive genes *CRYAB* and *NTRK2* are downregulated in the pTDPneg group, while another astrocyte-reactive gene, *KCNJ10*, is upregulated in the pTDPhigh group (**Fig. 5c**). These data suggest that there might be a positive correlation between astrocyte reactivity and cortical pTDP-43 accumulation. *NTRK2* encodes the neurotrophic tyrosine receptor kinase 2, TrkB, which interacts with brain-derived neurotrophic factor (BDNF), a crucial neurotrophin for brain function. BDNF regulates neuronal survival and synaptic plasticity. When TrkB is activated, it triggers a positive feedback loop and upregulates the transcription of BDNF through MAPK pathways^59^. One of the mechanisms by which astrocytes provide neurotrophic support is to release BDNF. Deprivation of BDNF and the TrkB signaling pathway increases inflammatory cytokines and promotes neuronal cell death^60^, leading to the suggestion that BDNF/TrkB deficiency contributes to AD pathogenesis. A decrease in TrkB expression is also found in the postmortem brains from AD patients^61^, suggesting a common dysregulation shared between *C9orf72* ALS/FTD and AD. This prior evidence together with our new findings suggest a potential impact of astrocytes on neuronal survival not just in late disease stages but also early before the accumulation of pTDP-43.

Several genes exhibited distinct expression changes specifically in the pTDPhigh group in astrocytes. For example, *SLC38A2*, encoding the protein SNAT2 primarily located in astrocyte processes^62^, was significantly upregulated in the pTDPhigh group. NEAT1 also showed elevated expression levels in astrocytes (**Fig. 5d**). NEAT1 expression was notably higher in non-neuronal cells compared to neurons (**Fig. 5e**). NEAT1 is a long non-coding RNA that has been proposed to function as a structural scaffold for assembling paraspeckles^63^, membraneless nuclear bodies composed of proteins and RNA. Many proteins present at paraspeckles are RNA-binding proteins involved in RNA splicing or post-transcriptional regulation and nuclear retention of RNAs, including TDP-43. NEAT1 is bound strongly by TDP-43^64^ and it has been shown that TDP-43 is involved in regulating the alternative polyadenylation switch of NEAT1^65^, resulting in altered levels of the NEAT1_1 short and NEAT1_2 long isoforms. Depletion of TDP-43 ALS/FTD in human embryonic stem cells results in increased levels of the NEAT1_2 long isoform, increased number of paraspeckles, and cell differentiation^65^. It has been shown that the G_4_C_2_ foci found in *C9orf72* ALS/FTD patient derived fibroblast colocalize with paraspeckle proteins but not NEAT1, suggesting that *C9orf72* G_4_C_2_ RNA could form a distinctive paraspeckle class that is independent of NEAT1^66^. However, it is unclear how NEAT1 expression and paraspeckle formation in astrocytes could be related to astrocyte reactivity, and whether this is due to the lack of nuclear TDP-43 or the increase of *C9orf72* toxic G_4_C_2_ RNAs. Previous studies have linked NEAT1 overexpression in the hippocampus of mice to impaired memory formation^67^. Dysregulation of NEAT1 and paraspeckle function has been observed in several neurodegenerative diseases^68^. Increased expression of NEAT1 in astrocytes of pTDPhigh samples suggests that astrocytes in late disease stage might be reactive due to dysregulation of RNA retention and/or number of paraspeckles.

We observed upregulation of *RPS6KA2*, which encodes RSK3, a ribosomal protein S6 kinase, in astrocytes of Emory pTDPhigh samples. We also detected upregulation of *RPS6KB1*, which encodes S6K1, another ribosomal protein S6 kinase, in the Mayo pTDPhigh samples (**Fig. 5f,g**). Although not statistically significant, we also noted elevated expression of *RPS6KA2* in excitatory neurons of pTDPhigh samples (**Fig. 5f,g; Supplementary Table 4b)**. Ribosomal protein S6 kinases, particularly S6K1, are well-known downstream effectors in the mTORC1 signaling pathway^69,70^, with recent studies indicating that mTORC1 activation promotes astrocyte development^71^. RSKs, including RSK3, target various substrates^72^, including Raptor, suggesting its involvement in mTORC1 signaling and its potential to regulate translation and cell survival by phosphorylating eukaryotic translation initiation factor-4B (eIF4B). To validate these results, we performed immunoblotting on protein lysates extracted from frontal cortical tissues of Emory cohort samples for phosphorylated S6 and total S6 protein. We observed significant increases in phosphorylated S6 levels specifically in pTDPhigh samples compared to controls (**Fig. 5h-i; Supplementary Fig. 7**), and reduced phosphorylated S6 in pTDPneg and pTDPmed samples, although this decrease is not statistically significant (**Fig. 5h,i**). Total S6 protein levels showed variation (**Fig. 5i**), suggesting a disease staging trend with higher levels in pTDPneg and lower levels in pTDPmed and pTDPhigh samples, indicating a decrease as pTDP-43 accumulation increases. Reduced RSK3 expression has been suggested to alleviate the neurodegenerative phenotype observed in Spinocerebellar Ataxia Type 1 (SCA1)^73^, another repeat expansion neurodegenerative disorder. RSK3 is highly expressed in the brain compared to other RSKs^74^ and possesses a potential nuclear localization signal (NLS). These findings collectively suggest differential activation or inactivation of signaling involving phosphorylated S6 protein in early and late disease stages of *C9orf72* ALS/FTD. The presence of DEGs shared between astrocytes and other cell types suggests the importance of astrocytic surveillance or homeostatic function in relation to surrounding cortical cells during neurodegeneration. In SOD1 ALS mouse models, downregulation of the SOD1 gene in astrocytes has been shown to slow disease progression^75^, providing further evidence of the crucial role of astrocytes in ALS/FTD disease progression.

### Heterogeneity of pTDP-43 accumulation in inhibitory and excitatory neurons

A total of 11,835 nuclei can be annotated as excitatory or inhibitory neurons using gene activity scores for key linage genes in the Emory cohort, and each category consists of 9 and 6 clusters, respectively (**Fig. 6a,b**). Thus, neurons are the most diverse cell type in the single nucleus multiome dataset in the Emory cohort. In contrast, control samples and *C9orf72* ALS/FTD samples with varying levels of pTDP-43 from the Mayo cohort have much fewer neuronal nuclei (**Supplementary Fig. 3**) and we have excluded the Mayo cohort from the following neuronal specific analysis. A similar issue has been observed by others when using *C9orf72* FTD frontal cortex samples from the Mayo Clinic Brain bank^22^. Multiple neuronal subtypes can be annotated based on known marker genes^76^. Excitatory neurons can be categorized by their cortical layer position (layer 2-6) and their axonal projections (**Fig. 6c**); while inhibitory interneurons can be grouped by their developmental origin from the medial, lateral or caudal ganglionic eminences and classified based on their subtypes (**Fig. 6d**). It is not known which neuronal cell types are more vulnerable to pTDP-43 accumulation and/or nuclear loss of TDP-43. NeuN+ cortical neurons from the neocortex of *C9orf72* ALS/FTD patients have been fractionated previously based on levels of nuclear TDP-43, allowing the characterization of nuclear TDP-43 positive and negative specific transcriptomes using bulk RNA sequencing^77^. However, this study was not able to identify the neuronal cell types contributing to the ensemble of TDP-43 positive and negative RNA-seq profiles. To estimate the contribution of neuronal subtypes present in this TDP-43 sorted neuronal population, we employed the cell composition deconvolution algorithm CIBERSORTx^78^ and compared our single nucleus datasets with the published TDP-43 sorted bulk transcriptomes on NeuN-positive nuclei. We were thus able to quantify the contribution of each individual neuronal subtype identified in our multiome dataset to the published TDP43-negative and TDP43-positive transcriptomes. It has been shown that NeuN positive neurons are typically composed of 70% excitatory neurons and 30% inhibitory neurons^76^, and we found our deconvolution analysis performs as expected in that more than 70% of NeuN-positive transcriptomes correspond to excitatory neuronal clusters (an average of 92.9% and 78.5% of cells in the TDP43-negative and TDP43-positive transcriptomes, respectively) (**Fig. 6e**). Among all neuronal clusters, EX-1, a cluster consisting of cortical projection neurons with high expression of CUX2 and LAMP5, has the most significant contribution to the nuclear TDP43-negative cells (**Fig. 6e**). This result suggests that a significant proportion of excitatory neurons with high expression of CUX2 and LAMP5 have nuclear TDP-43 loss, distinct from other neuronal populations.

**Fig. 6.**
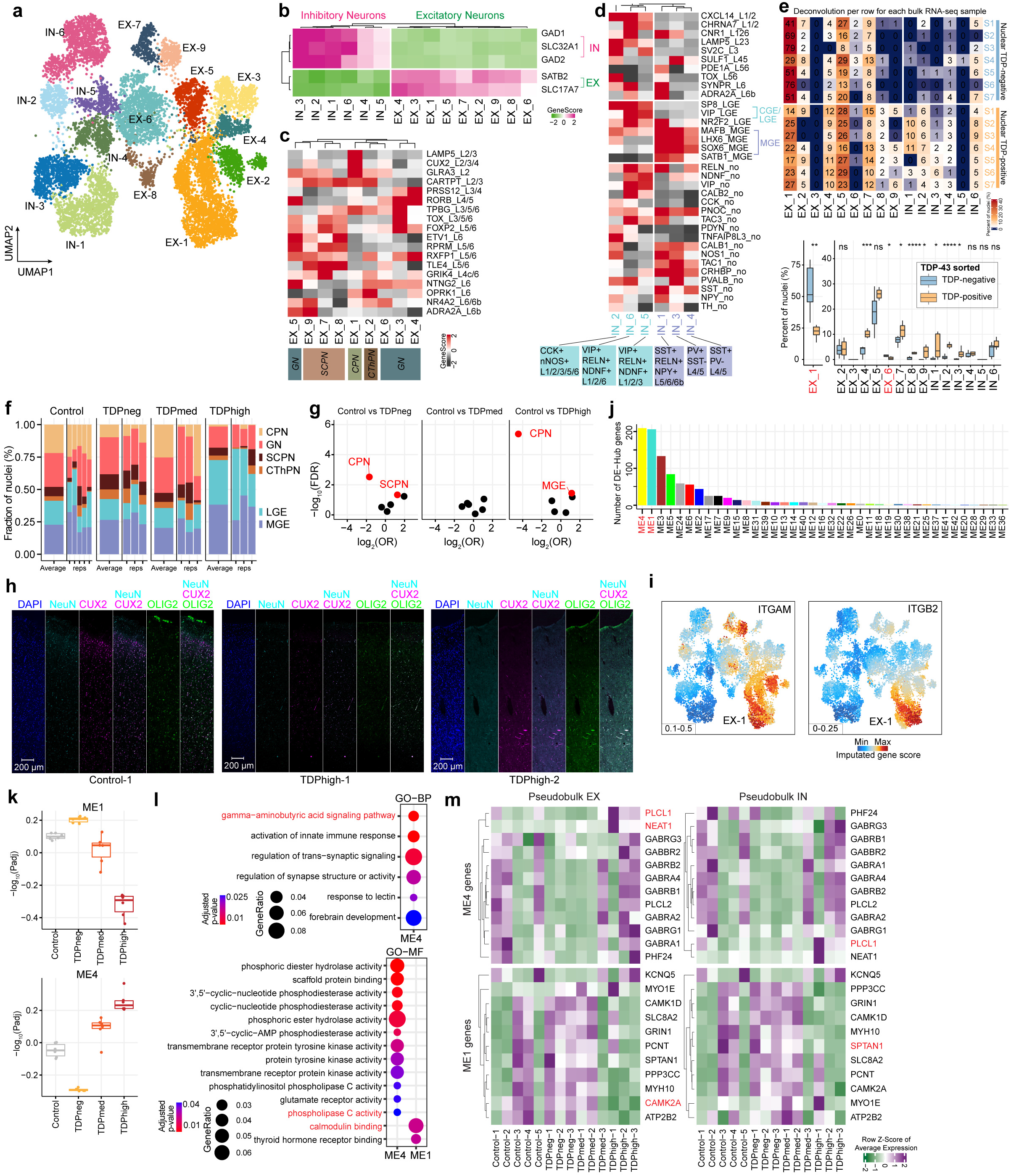
Neuronal cell types in the prefrontal cortex of control and *C9orf72* ALS/FTD donors. (a) UMAP plots of neuronal clusters. (b) Gene activity scores for marker genes of excitatory and inhibitory neurons. (c) Heatmap of gene activity scores of cortical layer specific marker genes for excitatory neurons. Axonal projection subclassification is indicated below. CPN, cortical projection neurons; GN, granule neurons; SCPN, subcortical projection neurons; CThPN, corticothalamic projection neurons. (d) Heatmap of gene activity scores of marker genes associated with inhibitory neurons of subpallial origin (top), cortical layers (middle) and subclassification (bottom). CGE, caudal ganglionic eminence; MGE, medial caudal ganglionic eminence; LGE, lateral ganglionic eminence; SST, somatostatin; RELN, reelin; NPY, neuropeptide Y; PV, parvalbumin; VIP, vasoactive intestinal peptide; NDNF, neuron-derived neurotrophic factor; CCK, cholecystokinin; nNOS, neuronal nitric oxide synthase. (e) Top: summary of cell proportion deconvolution with pTDP-43 positive and negative nuclei (n.s. not statistically significant; P≤0.05 is considered statistically significant: * P≤0.05, ** P≤0.01, *** P≤0.001, **** P≤0.0001); bottom: heatmap representation of cell proportion deconvolution data in each individual nuclear pTDP-43 positive and negative transcriptome. (f) Proportion of neuronal subtypes defined by cortical projection or developmental origins in all sample groups. (g) Volcano plots showing odds ratio (OR) and FDR computed by MASC^103^ for all the neuronal subtypes. Red labeled neuronal subtypes that are significantly increased or depleted in association with specific *C9orf72* ALS/FTD donor groups (FDR-adjusted P < 0.05; absolute OR >0). (h) Immunostaining of human postmortem cortical tissue for the pan-neuronal marker NeuN (cyan), oligodendrocyte lineage marker OLIG2 (green), CUX2 (magenta) and DAPI in blue. (i) Gene activity score for the *ITGAM* and *ITGB2* genes that encode the heterodimer C3 receptor. (j) Number of differential hub genes found in each module. (k) Significance of WGCNA modules with different levels of pTDP-43. (l) Gene ontology analysis of the differential hub genes in the ME1 and ME4 modules. (m) Heatmap demonstrating the average gene expression of identified hub genes in modules ME1 and ME4 across all samples from the Emory cohort.

### CUX2+ cortical projection excitatory neurons are significantly reduced in the frontal cortex of *C9orf72* ALS/FTD donors

ALS/FTD with pTDP-43 inclusions is typically accompanied by frontal cortex atrophy and neuronal loss^9^. We therefore grouped neurons based on their excitatory projection classification and developmental origin for interneurons to avoid cell clusters with few nuclei. We found that the proportion of cortical projection neurons is more than three-fold lower in TDPhigh and TDPneg patient groups compared to control (**Fig. 6f**), suggesting that these neurons are especially susceptible to *C9orf72* ALS/FTD frontal cortex degeneration regardless of the frontal cortical pTDP-43 levels. We systematically assessed the differential abundance between *C9orf72* ALS/FTD donor and control groups for all neuronal clusters. Cortical projection neurons showed significant proportional changes in both TDPhigh and TDPneg donor groups, while subcortical projection neurons and inhibitory neurons originating from the medial caudal ganglionic eminence showed significant proportional changes in pTDPneg and pTDPhigh groups, respectively (FDR-adjusted P < 0.05, absolute log2(odds ratio (OR)) > 0; Methods; **Fig. 6g**). To confirm the loss of CUX2 neurons in the upper cortical layers, we used immunofluorescence microscopy using antibodies to CUX2 and cell-type specific marker proteins in pTDPhigh samples (**Fig. 6h**). This result also further confirms our previous conclusion on the vulnerability of cortical projection neurons in *C9orf72* ALS/FTD degeneration, despite the differences in pTDP-43 accumulation. Also, loss of cortical projection neurons in TDPhigh donors is more extensive than in TDPneg donors (**Fig. 6f,g**). We found that the genes *ITGAM* and *ITGB2*, which encode for subunits of the CR3 complement factor C3 receptor, are specifically highly expressed in cortical projection neurons compared to other neuron types (**Fig. 6i**). This finding supports our earlier speculation that cortical projection neurons might be specifically tagged by C3 released by microglia in the early stages of disease, leading to microglia mediated synapse loss and phagocytosis of neurons. This novel finding could also explain the specific vulnerability of cortical projection neurons in early disease stages of *C9orf72* ALS/FTD.

In addition to cortical projection neurons, inhibitory neuronal clusters originating from the medial caudal ganglionic eminence (MEG) are significantly increased in proportion in the TDPhigh donor group compared to control (**Fig. 6f,g**). Based on our cell composition deconvolution analysis against pTDP-43 positive and negative specific transcriptomes, IN-1 and IN-3 MEG originated neurons have a higher contribution to the pTDP-43 positive compared to the pTDP43-negative transcriptome (**Fig. 6e**). The results suggest that inhibitory neurons originated from the MEG might be resistant to neurodegeneration, possibly because they are less vulnerable to nuclear TDP-43 loss. Interestingly, the TDPmed donor group does not have significant changes in the relative proportion of neuronal clusters, perhaps because this group represents a transitional state in disease progression that reflects a mixture of cortical neuronal populations.

Since neurons exhibit many differentially expressed genes in all *C9orf72* ALS/FTD donor groups, we set out to identify putative gene regulatory networks that correlate with pTDP-43 accumulation. We used the weighted gene co-expression network analysis (WGCNA) to cluster co-expressed genes found in neurons into modules and to identify highly correlated genes. We identified 43 modules (**Supplementary Fig. S8a**) and among these modules, two of them, ME1 and ME4, significantly correlate with pTDP-43 levels (**Fig. 6j,k and Supplementary Fig. S8b**). ME4 positively correlates while ME1 negatively correlates with the amount of pTDP-43 (**Fig. 6k**). We found these two modules also have the most differentially expressed hub genes (**Fig. 6l and Supplementary Table 7a**), further suggesting that the differentially expressed hub genes in these two modules are strongly associated with disease progressive changes in neurons. Based on gene ontology analysis of these differential hub-genes in the ME4 and ME1 modules, we found enrichment of distinctive set of genes. For example, genes involved in the gamma-aminobutyric acid (GABA) signaling pathway are enriched in the ME4 module whereas genes involved in calmodulin binding are enriched in the ME1 module (**Fig. 6l and Supplementary Table 7b)**. Specifically, *PLCL1* is found in multiple GO enriched terms, including modulation of the GABA signaling pathway, inositol lipid-mediated signaling, and phospholipase C activity (**Supplementary Table 7b**). *PLCL1* encodes for phospholipase C-related inactive protein type 1, and it is significantly upregulated in pTDPhigh samples in both excitatory and inhibitory neurons (**Fig. 6m**). PLCL1 has been shown to regulate GABA receptor trafficking and it has been found to be upregulated in the dopaminergic neurons in the substantia nigra^79^, suggesting this might be another common link to other neurodegenerative disorders outside of the frontal cortex. *NEAT1* emerges as another prominent hub gene in the ME4 module (**Fig. 6m**), showing specific upregulation during the late disease stage in pTDPhigh samples. As previously discussed, *NEAT1* typically exhibits high expression in glial cells and low expression in neurons. This finding further suggests that dysregulation of paraspeckles in excitatory neurons could be another characteristic feature of the late disease stage with high levels of cortical pTDP-43. In contrast to ME4 hub genes, *SPTAN1* represents a strong ME1 module hub gene that is upregulated in early disease stages. *SPTAN1* encodes a spectrin family protein, a crucial component of the cytoskeleton^80^. Notably, it has been demonstrated to interact with calmodulin and participate in calcium signaling. Pathogenic variants in spectrin genes are implicated in various neurological disorders, including cerebellar ataxia. Furthermore, *CAMK2A*, another differential hub gene in the ME1 module, exhibits specific downregulation in the pTDPhigh group. Exon skipping of CAMK2A transcripts has been observed in TDP-43 knockdown mouse primary neurons^81^, suggesting that the downregulation of *CAMK2A* could be a direct consequence of the loss of nuclear TDP-43 in pTDPhigh samples. Additionally, *CAMK2A* has been reported to have reduced protein abundance in the cerebrospinal fluid of ALS patients compared to controls^82^. This comprehensive WGCNA analysis in neurons further bolsters our earlier findings, indicating that a distinct set of genes is involved in early and late disease stages in a cell-type-specific manner.

## Discussion

A G_4_C_2_ repeat expansion in the first intron of the *C9orf72* gene is the most common genetic cause of both FTD and ALS. Neuromuscular abnormalities observed in patients with ALS are caused by degeneration of motor neurons in the motor cortex of ALS/FTD patients^83^. However, the contribution of different cell types of the frontotemporal cortex to neurodegeneration and cognitive decline during FTD disease progression and the accompanying molecular changes remain largely unexplored. In this study, we utilized a unique *C9orf72* ALS/FTD staging paradigm by selecting cases based on the abundance of pTDP-43. Cortical cytoplasmic accumulation of pTDP-43 has been found to correlate with neuropathological burden and severity of FTD clinical symptoms, and the progression of pTDP-43 distribution in the CNS has been proposed to stage patients in different phases of the disease^84^. We utilized a multiome approach to simultaneously analyze changes in chromatin accessibility and gene expression in the same cell. Using this approach, we identified several systematic changes in the early and late stages of disease not previously reported. These include the loss of neurosurveillance microglia, significant increases in phosphorylated ribosomal S6 protein, and global dysregulation of chromatin accessibility uniquely found in non-neuronal cells associated with high pTDP-43 levels. We also observed abnormalities in oligodendrocytes, microglia, and astrocytes specifically associated with late disease stages. Interestingly, cortical projection neurons appear to be selectively vulnerable to *C9orf72* ALS/FTD progression.

We observed more pronounced disease stage specific changes in glial cells compared to neurons. Changes observed in *C9orf72* ALS/FTD donors can be interpreted as being a consequence of this gene, presence of repeats in the RNA, or the presence of peptides translated from these repeats. Our findings suggest that microglia may be the first cells to respond to miss expression of *C9orf72* in the frontotemporal cortex in the initial stages of FTD disease, before aggregation of pTDP-43 in the cytoplasm, by activating the complement cascade and increasing phagocytosis as well as by altering their neuronal surveillance activity. This pattern mirrors observations in Alzheimer’s disease, albeit with different genetic contributions, suggesting a common activation of the microglia immune response in early neurodegeneration in both diseases. However, in late disease stages, the interferon response is activated in microglia instead. Astrocytes exhibit alterations both early and late in disease progression, including the downregulation of astrocyte-reactive genes like *CRYAB* and *NTRK2* before the formation of pTDP-43 inclusions. During late disease stages, astrocytes significantly upregulate *NEAT1* and *RPS6KA2*. This raises the question of the role of paraspeckles in in late-stage pathology, since TDP-43 protein is known to bind to NEAT1 and colocalize in paraspeckles with other splicing regulatory proteins^65^. The increased *RPS6KA2* gene expression and phosphorylated ribosomal protein S6 levels specifically in astrocytes during late disease stages are particularly intriguing. Since *RPS6KA2* encodes RSK3, the only RSK with a potential nuclear localization signal, it is possible that upregulation of *RPS6KA2* may lead to the phosphorylation of additional nuclear proteins with roles in the regulation of gene expression. Furthermore, the upregulation of both *RPS6KA2* and *NEAT1* in astrocytes and neurons suggests a shared molecular mechanism between these cell types, potentially expanding our current understanding of their interactions.

The most significant finding unique to donors in late disease stages with high levels of pTDP-43 is the high proportion of newly differentiated/premyelinating oligodendrocytes (ODC-2), which is not observed in control or *C9orf72* ALS/FTD samples with low pTDP-43 accumulation. Dysregulation of oligodendrocyte maturation and function may be a direct consequence of the formation of pTDP-43 cytoplasmic inclusions or the nuclear loss of TDP-43. Indeed, pTDP-43 inclusions in oligodendrocytes are a hallmark of *C9orf72* ALS/FTD. The relatively low expression of genes encoding myelin protein components in this cell cluster may be due to high cytoplasmic pTDP-43 accumulation, since nuclear TDP-43 binds to transcripts encoding for myelin proteins^32^ and soluble cytoplasmic TDP-43 is involved in the posttranscriptional regulation of myelin proteins^31,33^. We do not observe a significant change in oligodendrocyte progenitor cells, suggesting that the cluster of premature oligodendrocytes is likely a result of its inability to become mature due to the downregulation of myelin components and failed to undergo the typical programmed cell death observed for the majority of premyelinating oligodendrocytes during adult oligodendrogenesis. Several lines of evidence converge on oligodendrocyte dysfunction as an important contributor to ALS/FTD pathogenesis^25,85–87^. Tissues from sporadic ALS patients show significant regions of demyelination and decreased expression of myelin related proteins^25^. Genetic analyses have also provided insights into the role of oligodendrocytes in ALS/FTD. Recent GWAS studies have implicated single nucleotide polymorphisms (SNPs) in the *MOBP* gene, which encodes for myelin-associated oligodendrocyte basic protein, as a risk factor for ALS^88,89^. SNPs in *MOBP* are also associated with shorter disease duration and more severe white matter degeneration in FTD^90^. It is thus possible that the cause of cortical projection neuron loss in TDPhigh donors is the lack of sufficient mature oligodendrocytes, which are essential for neuron myelination and metabolic support. The impairment of myelination is not limited to *C9orf72* FTD TDP-43 pathology. In AD donors carrying two copies of the APOE4 variant, cholesterol homeostasis is responsible for the downregulation of myelin-associated genes in oligodendrocytes^91^. However, the downregulation of myelin-associated genes in *C9orf72* TDPhigh oligodendrocytes is not accompanied by changes in cholesterol homeostasis, suggesting that cortical myelination defects are common in patients with cognitive impairments with different genetic mutations. Further dissection of oligodendrocyte-neuron interactions may give additional insights into the mechanisms underlying the progression of FTD. The CUX2+ cortical projection neurons are lost both in pTDPneg and pTDPhigh samples, suggesting neuronal loss takes place early in disease progression. However, we found changes in gene expression in neurons unique to early and late disease stages. It is possible that these changes in the neuronal transcriptome can result in neuronal loss or that changes in the neuronal transcriptome are a consequence of dysregulation of other cell types. One possibility is that, since the cortical projection neurons specifically express high levels of C3 receptor subunits, it is possible that synapse pruning and phagocytosis are mediated by C3 released by the complement activated microglia. While our study comprises a relatively small number of samples, the approach of grouping samples based on cortical pTDP-43 levels allowed us to distinctly map out novel disease progression events and to map new and previously reporter findings to specific disease stages. It will be important to explore whether these findings extend to larger cohort studies. Ultimately, the systematic identification of cell-type-specific defects in pathways common to all *C9orf72* ALS/FTD donors, as well as disease stage-specific alterations, will inform the targets and timing of therapeutic interventions.“

## Methods

### Human tissue samples

Post-mortem brain samples from the dorsolateral prefrontal cortex (DLPFC, Brodmann area 9; BA9) of *C9orf72* ALS/FTD patients and controls were obtained from the Goizueta Emory Alzheimer’s Disease Center Brain Bank and Mayo Clinic Brain Bank with approval from the respective Institutional Review Board. All *C9orf72* patients had a clinical diagnosis of ALS and/or FTD. Controls consisted of normal individuals with no clinical history of neurological disease. Patient information is provided in **Supplementary Table 1**. All brains underwent thorough neuropathologic evaluation, including hematoxylin and eosin stains, silver stains, and immunohistochemistry for β-amyloid, tau, α-synuclein, and phosphorylated TDP-43. Repeat primed PCR was performed on all samples to confirm the presence of expanded repeats in the *C9orf72* locus (**Supplementary Table 1**). Immunohistochemistry and quantitative immunoassay measurements for dipeptide repeat proteins were also performed on all Emory cases as an alternative method to confirm the *C9orf72* repeat expansion (**Supplementary Table 1**).

### Quantification of cortical phosphorylated TDP-43 levels

Sequential biochemical fractionation was performed first and followed by Meso-Scale Discovery (MSD) immunoassay. Tissue lysates were fractionated according to previously published protocols^9^. In brief, ∼200 mg of dorsolateral prefrontal cortex (DLPFC) tissue was homogenized in low salt buffer (10 mM Tris pH 7.5, 5 mM EDTA pH 8.0, 1 mM DTT, 10% sucrose, 1x HALT protease/phosphatase inhibitors). Lysates were pelleted at 25,000 g for 30 min. Supernatants were collected as the “low salt” fraction. The resulting pellet was solubilized in Triton-X buffer (1% triton X-100 and 0.5 M NaCl in low salt buffer). Lysates were subsequently pelleted at 180,000 g for 30 min. Supernatants were collected as the “Triton-X” fraction. The resulting pellet was solubilized in Triton-X buffer with 30% sucrose. Lysates were subsequently pelleted at 180,000 g for 30 min. The resulting pellet was solubilized in Sarkosyl buffer (1% sarkosyl and 0.5 M NaCl in low salt buffer). Lysates were then pelleted at 180,000 g for 30 min. Supernatants from these fractions were not used for analysis. The resulting insoluble pellet was resolubilized in 8 M urea solution (pH 8.0) and used for MSD immunoassay to measure blinded the abundance of phosphorylated TDP-43 in the detergent insoluble and urea soluble fractions using a previously described sandwich immunoassay that utilizes MSD electrochemiluminescence detection technology^92^. The capture antibody was a mouse monoclonal antibody that detects TDP-43 phosphorylated at serines 409/410 (1:500, no. CAC-TIP-PTD-M01, Cosmo Bio USA, was used for Emory samples; 2 μg/mL, no. 22309-1-AP, ProteinTech, was used for Mayo samples), and the detection antibody was a sulfo-tagged rabbit polyclonal C-terminal TDP-43 antibody (2 μg/mL, 12892-1-AP, Proteintech). Lysates were diluted in 8 M urea solution (pH 8.0) such that all samples of a given type were made up to the same concentration and an equal amount of protein for samples was tested in duplicate wells. Response values corresponding to the intensity of emitted light upon electrochemical stimulation of the assay plate using the MSD QUICKPLEX SQ120 were acquired. These response values were background corrected by subtracting the average response values from corresponding negative controls e.g., lysates from tissues or cells lacking a repeat expansion per batch.

### Immunohistochemistry

Immunohistochemistry for pTDP-43 was done as previously described^93^. In brief, paraffin embedded sections (8 µm) were deparaffinized in Histo-clear (National Diagnostics) and rehydrated in 100% and 95% ethanol, followed by water. Steam heat antigen retrieval was performed for 30 min. To prevent non-specific chromogen development, we quenched endogenous peroxidase activity using hydrogen peroxide followed by 3x washes in TBS-T (Tris buffered saline solution with 0.05% Tween-20). Tissue sections were blocked using serum-free protein block (Dako) for 1 h. Primary antibodies to pTDP-43 (Cosmo Bio USA, TIP-PTD-P02) were applied for 45 min at room temperature, followed by 3x washes in TBS-T. Polymer HRP-conjugated secondary antibodies (Dako) were applied for 30 min at room temperature. Peroxidase labeling was visualized with 3,30 -diaminobenzidine (DAB). Sections were counterstained with Gill’s hematoxylin and Scott’s tap water substitute was used as the bluing reagent.

#### Isolation of nuclei from frozen brain tissue

Tissue sections were snap frozen according to each brain bank’s specification and stored at −80°C and the nuclei were isolated as previously described^94,95^. Briefly, 20 mg frozen tissues were thawed in 1 mL cold homogenization buffer (260 mM sucrose, 30 mM KCl, 10 mM NaCl, 20 mM Tricine-KOH pH 7.8, 1 mM DTT, 0.5 mM Spermidine, 0.2 mM Spermine, 0.3% NP40, cOmplete Protease inhibitor (Roche), and Ribolock) and homogenized in a pre-chilled Dounce. Cell lysates were passed through a 70 µm Flowmi cell strainer before separation using a discontinuous iodixanol gradient and centrifugation at 1480 g at 4°C for 20 min in a swinging bucket centrifuge with the brake off. The nuclei band located at the interface between 30% and 40% iodixanol was collected and washed in RSB-T wash buffer (10 mM Tris-HCl pH 7.4, 10 mM NaCl, 3 mM MgCl_2_, 0.1% Tween-20).

#### Single-nucleus multiome library preparation and sequencing

Libraries were generated using the 10x Genomics Chromium Single Cell Multiome ATAC + Gene Expression kit following the manufacturer’s instructions, with the following modifications. Per sample, 16,100 nuclei were resuspended in 1x diluted nuclei buffer (10x Genomics) with 2% BSA (Sigma) with a capture target of 10,000 nuclei. First, ATAC-seq libraries were sequenced to target of 25,000 read-pairs per nucleus and RNA libraries were sequenced to 20,000 read-pairs per nucleus on an Illumina NovaSeq 6000 instrument at the Florida State University Translational Science Laboratory and NovaSeq X Plus instrument at Admera Health. The matched RNA-seq and ATAC-seq libraries were processed using the 10x Genomics Cell Ranger ARC (cellranger-arc-2.0.0) pipeline with default parameters and aligned to the hg38 human genome assembly (refdata-cellranger-arc-GRCh38-2020-A-2.0.0 We aim for 50% saturation in snRNA-seq and 30% saturation in snATAC-seq libraries for each sample. Supplementary Table S2 contains the summarized sample matrix generated by cellranger-arc count, encompassing all nuclei information before stringent filtering. The subsequent files produced by cellranger-arc count, including the filtered gene-barcode matrices for snRNA-seq (filtered_feature_bc_matrix.h5) and deduplicated ATAC-seq fragment files (atac_fragments.tsv.gz), were utilized for further processing and quality control analysis.

#### Processing and analyses of single nucleus multiome data

Reads mapping to the mitochondrial genome, chromosome Y, and common blacklisted regions were excluded from downstream analysis for both snATAC-seq and snRNA-seq libraries. ArchR (v1.0.2)^17^ and Seurat (v4.1.0)^18^ were used for processing the paired snATAC-seq fragment data and snRNA-seq gene expression data for each sample. To ensure that only high-quality nuclei proceeded to downstream analysis, stringent quality control filtering steps were implemented using paired snRNA-seq and snATAC-seq libraries for each sample. Nuclei were excluded from downstream analysis if they met the following criteria: a snATAC-seq TSS score <4, fewer than 1000 unique nuclear fragments in snATAC-seq, and lack of matched RNA reads. Nuclei doublets were excluded using ArchR addDoubletScores() and filterDoublets() functions using snATAC-seq TileMatrix for each sample. Moreover, for snRNA-seq quality control, only nuclei meeting the following criteria in the RNA quality control matrix were retained: a log10(number of genes per UMI) greater than 0.8, mitochondrial RNA levels less than 1%, and a gene count ranging from 500 to 5000 for the Emory samples and from 200 to 10000 for the Mayo samples. Only the nuclei that passed quality control were used for downstream analyses. An average of 4.6 % nuclei doublets were removed using the ArchR doublet detection tool and default parameters. As a result of the stringent quality control, a total of 34,874 nuclei were used for downstream analysis with median TSS score of 9.895 and median fragments per nucleus of 12068 for Emory samples. A total of 53,331 nuclei with a median TSS score of 7.318 and median fragments per nucleus of 9805 for samples of the Mayo cohort were used for downstream analysis. The quality control matrix is provided in **Supplementary Fig. 1 and Supplementary Table 2**. We performed pseudobulk per sample correlation analysis and PCA analysis of pseudo-snRNA-seq data for both cohorts (**Supplementary Fig. 4b,c**) and found a strong batch effect between Emory and Mayo samples, however, all samples shows strong correlations within each sample group. This information suggests the presence of batch effects based on the brain bank of origin. We also conducted additional analyses to correlate marker genes across both datasets, revealing a high degree of similarity, as depicted in **Supplementary Fig. 4d**. Notably, oligodendrocytes exhibited the highest similarity in marker genes between the two datasets, providing further evidence for the presence of common oligodendrocyte dysfunction in both cohorts. Because there is a strong batch effect between Emory and Mayo samples, the two cohorts were processed independently and in parallel for all downstream analyses.

We first performed dimensionality reduction using each GeneExpressionMatrix derived from snRNA-seq and TileMatrix derived from snATAC-seq separately for each sample with default parameters using ArchR’s addIterativeLSI() on each reduced dimension, followed by Uniform Manifold Approximation and Projection. We then projected both datasets and observed a strong sample batch effect in the snRNA-seq data; however, this effect was not observed in the paired snATAC-seq libraries (**Supplementary Fig. 2a,b**). To further assess the potential impact of sample preparation batch effects on the GeneExpression Matrix of the snRNA-seq data, we imported the gene-UMI count matrix from nuclei that passed the quality control in the preprocessing step described above from all samples into Seurat (v4.1.0)^18^. We then conducted pseudobulk analysis. This involved aggregating the UMI counts of genes mapped to autosomes and chromosome X from each sample. We then performed regularized log transformation, normalizing with respect to library size using the DESeq2 rlog() function (v1.34.0)^23^, followed by principal component analysis. The data were labeled according to sample identity, levels of pTDP-43, sex, and sample library preparation batch (**Supplementary Fig. 2d**). We did not observe any specific batch effect related to sample or pTDP-43 grouping at both PC1 and PC2. However, we noticed a slight variation in sex-specific differences at PC2 (8%). Nevertheless, a pronounced batch effect stemming from sample library preparation was evident. To address this, we employed two types of normalization methods known for correcting batch effects. For size-factor normalization, we utilized Seurat’s NormalizeData() function, setting normalization.method to ‘RC’ and scale.factor to 1e6. For sctransform normalization, we utilized Seurat’s SCTransform() function, using mitochondrial mapping percentage and sample library preparation as variables for regression. Both normalization methods effectively mitigated the batch effect. Therefore, these covariates were taken into account during the performance of the differential gene expression analysis. Consequently, we also applied Harmony batch correction^19^ on the snRNA-seq and snATAC-seq datasets. This correction incorporated sample, groups categorized by pTDP-43 levels, and sample preparation batch as covariates to mitigate the batch effects. This process effectively removed the batch effect in both datasets. Subsequently, we combined the reduced dimensions of both datasets using ArchR’s addCombinedDims() before and after applying Harmony batch correction, confirming the successful removal of the batch effect from the libraries (**Supplementary Fig. 2c**)

### Identification of cluster and cell type assignments

Cell clusters were called using Seurat implemented in ArchR using the combined single nucleus RNA and ATAC matrix with a resolution setting of 1.2 and 0.8 for Emory and Mayo samples, respectively (**Supplementary Fig. 3a**). Clusters containing less than 100 nuclei were excluded from subsequent analysis. In total, 41 and 20 cell clusters were distinguished for Emory and Mayo samples, respectively, all of which possessed known cortical cell type identities (**Supplementary Fig. 3a**). Cell type identification was performed based on gene activity scores calculated using ArchR with default parameters; the gene activity scores are correlated with gene expression and calculated based on chromatin accessibility at the gene body, promoter and distal regulatory regions^17,96^. Marker genes for each cluster were identified using ArchR getMarkerFeatures() function (filtering threshold: FDR ≤ 0.01 & log2(Fold change) ≥ 0.5; **Supplementary Table 3**) and manually compared to known marker genes of cortical cell types. The cell classification was further verified by gene modules computed using ArchR addModuleScore() function with geneScoreMatrix with the following genes for each cell type (**Supplementary Fig. 3d**): Neurons: *SNAP25* and *SYT1*; excitatory neurons: *SLC17A7*, *SATB2*, *RORB*, *NEUROD2*; inhibitory neurons: *GAD1*, *GAD2*, *NXPH1*; astrocytes: *GFAP*, *AQP4*, *SLC1A2*; microglia: *CSF1R*, *CD74*, *P2RY12*, *PTPRC*,*TMEM119*; oligodendrocytes: *MOBP*, *MBP*, *ST18*, *KLK6*, *SLC5A11*; oligodendrocyte precursor cells: *PDGFRA*, *CSPG4*; and endothelial cells: *FLT1*, *CLDN5*, *ABCB1*, *EBF1*. However, many of the Mayo samples have low number of neurons regardless of the levels of pTDP-43. Therefore, our main analysis framework focused on the Emory samples and we used the pseudobulk per major cell types found in the Mayo samples to validate our findings from the Emory cohort. In the Emory cohort, major cell types underwent additional subclustering analysis, excluding endothelial cells. This analysis included neurons, astrocytes, microglia, and oligodendrocyte lineage cells. Cluster identity analysis followed the previously described method. Subclusters exhibiting similar marker genes were merged. This yielded eight excitatory neuron clusters, six inhibitory neuron clusters, four astrocyte clusters, four microglia clusters, and seven oligodendrocyte lineage cell clusters. Following these two steps of cluster calling and identification, a total of 31 cell type-specific clusters were found in the Emory dataset.

### snATAC-seq peak calling and differential snATAC-seq chromatin accessibility analysis

ArchR was used to call peaks with default parameters in Emory samples. Since the clusters were less distinct in the Mayo cohort, the snATAC-seq peak calling and differential chromatin accessibility analysis were only performed in the Emory samples, not the Mayo samples. Briefly, a pseudo-bulk dataset was created for each of the major cell type using ArchR’s addGroupCoverages() function and the reproducible peak sets were called using addReproduciblePeakSet() with MACS2^97^ with a fixed-width peak size of 501 bp and iterative overlap peak merging based on coverage data grouped by each major cell type. The resulting PeakMatrix, with a total of 404,124 peaks, was used for downstream analysis. These peaks were designated as chromatin accessible regions. To identify differential chromatin accessible regions, we extracted the pseudo-bulk number of insertions observed per cell in each major cell type for both control and *C9orf72* ALS/FTD samples from the ArchR project using the getGroupSE() with the Peak Matrix. Differential peaks within each major cell type were identified by comparing control samples to *C9orf72* ALS/FTD samples with varying levels of pTDP-43, utilizing DESeq2^23^ with multi-factor designs. We accounted for sex and sample preparation batch as fixed effect covariates. Only AR containing at least 30 fragments were included in the comparison^23^. The DARs are considered significantly different if they have an FDR-corrected p-value < 0.05 and an absolute log2(fold change) > 1 relative to the control group.

### Comparison of snRNA-seq differential gene expression

We performed differential expression analysis using the model-based method MAST with a linear mixed effect hurdle model^24^ to account for covariates that contributed to the batch effect described above. For each sample from the Emory cohort of *C9orf72* ALS/FTD donors, pseudo-bulk data from each distinct cluster and six major cell types were compared to that from the healthy control samples. The raw count is stored in the “RNA@counts” slots of the Seurat R object. Normalization to the sequencing depth, referred to as counts per million (cpm), was performed using Seurat’s NormalizeData() function by specifying normalization.method=”RC”, scale.factor=1e6; the cpm is stored in the “RNA@data” slots of the Seurat R object. We also performed the independently SCTransform normalization to account for the confounding sources of variation of mitochondrial mapping percentage and sample preparation batch effect, and this data is stored in in the “SCT@counts” slots of the Seurat R object. The Seurat R object was converted to SingleCellExperiment object, and log2(count+1) was used to run MAST. Genes that were not expressed in at least 10% of nuclei were excluded from differential comparisons. The following linear mixed model was utilized with MAST where x is log2-normalized gene expression; *T* is the pTDP-43 grouping; *G* is the number of genes detected per nucleus, *U* is number of UMI, *M* is the percentage of reads mapped to the mitochondrial genome, *A* is the age of the subject, and these covariates were centered and scaled; *E* is the sex of the subject and *S* is the ID of the subject.

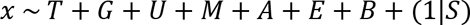

All terms were treated as fixed-effect terms except for the subject, which was treated as a random effect term. Differential expression of genes (DEGs) was identified using a likelihood ratio test (LRT), comparing the model between control and each pTDP-43 group. Hurdle p-values were generated by MAST, with p-values adjusted for multiple comparisons using the Benjamini & Hochberg false discovery rate (FDR) method. Additionally, fold changes (FC) were reported by the MAST model. To facilitate interpretation, counts-per-million fold changes (CPM FC) were computed by subtracting the mean log2CPM of selected nuclei in the control sample from the mean log2CPM of nuclei in the *C9orf72* samples. Genes were considered differentially expressed based on the following criteria: FDR < 0.05; model fold change > 1.1 and log2CPM > 1.5; convergence between model FC and average FC, with the difference between log2(model FC) and log2(CPM FC) < 2.

For each sample from the Mayo cohort, only pseudo-bulk data from seven major cell types were compared because many samples from the Mayo brain bank showed depletion of neuronal nuclei, as reported recently by others^22^. We have utilized MAST^24^, that allow us to model cells individually using a hurdle model. Because of low sample numbers, we did not employ the full random effect model as we did for the Emory samples. Instead, we have considered pTDP-43 grouping and number of genes detected per nucleus as fixed-effect terms. Differential gene expression results from both analyses were considered significant differentially if they had an FDR-corrected p-value < 0.05 and an absolute log2(fold change) > 0.5 relative to the control group.

#### Enrichment of TF motifs in differentially accessible regions

The regions identified as differentially accessible were tested for motif enrichment using ArchR peakAnnoEnrichment() after motif annotation analysis with addMotifAnnotations() using the cisbp motifs database^98^. The TF motifs were considered significantly enriched if FDR ≤ 0.1 & Log2FC ≥ 1.

### Immunostaining, imaging and quantifications using MERSCOPE

Frozen human brain tissue (prefrontal cortex, BA9 region) was sectioned at 10 µm thickness using a cryostat after embedding in O.C.T. compound (ThermoFisher Scientific Waltham, MA, USA). The MERSCOPE protein stain verification protocol (Vizgen, 10400112) was employed following the manufacturer’s instructions with the mouse, rabbit, goat protein stain verification kit. Briefly, sections were fixed in 4% paraformaldehyde and permeabilized in 70% ethanol for 24 hours at 4°C. Primary antibodies against FOX3 (NeuN, BioLegend, 834501, 1:1000), OLIG2 (R&D, AF2418, 1:20), and TCF7L2 (Cell Signaling, 2569, 1:1000) were used. Secondary staining solutions for anti-Mouse Aux4, anti-rabbit Aux5, and anti-goat Aux6 were applied, followed by gel embedding and incubation in a proteinase K-containing clearing solution. Sections underwent photobleaching for three hours using the MERSCOPE Photobleacher (Vizgen), followed by clearing at 37°C for 24 hours before imaging with the MERSCOPE (v233.230615.567) using protein stain verification reagent. Output VZG files were visualized for mosaic images using MERSCOPE visualizer (v2.3.3330.0). Output TIFF files were utilized for nucleus segmentation and staining intensity quantification. The raw DAPI staining image was stored as a 2D matrix, with each entry representing staining intensity. The matrix values were rescaled to range from 0 to 255, and a threshold of 80 was applied to generate a binary image indicating nucleus presence. Connected component detection using the Spaghetti algorithm identified nuclei, with area filtering (20 to 500 pixels) applied. For OLIG2, NeuN, and TCF7L2 protein staining images, intensities were rescaled to a range of 0 to 1000 for standardization. Within each nucleus, pixels from the protein staining image were extracted and mean intensity calculated as an estimate of protein abundance. The standard deviation of intensities within a cell was recorded as a measure of variation. Nuclei were classified as neurons if NeuN quantile was greater than 8 and OLIG2 quantile less than 3; nuclei with OLIG2 quantile greater than 8 and NeuN quantile less than 3 were classified as oligodendrocytes; all other nuclei were categorized as “other types”. TCF7L2 mean intensity was normalized by DAPI mean intensity to adjust for staining intensity variation. One-way ANOVA was utilized to compare significance between *C9orf72* ALS/FTD and control samples.

### Immunostaining and imaging using confocal microscopy

Frozen human brain tissue (prefrontal cortex, BA9 region) was sectioned at 40 µm thickness using a frozen vibratome. The sections were blocked for 1 h at room temperature (RT) in blocking buffer (PBS containing 5% bovine serum albumin, 1% normal donkey serum and 0.3% Triton X-100) and incubated in primary antibodies against FOX3 (NeuN, BioLegend, 834501, 1:1000), OLIG2 (R&D, AF2418, 1:20), TCF7L2 (Cell Signaling, 2569, 1:1000) and/or CUX2 (Proteintech, 82933-1-RR, 1:200) in blocking buffer for 24 at 4°C. The sections were washed three times for 5 min each at RT in PBS and then incubated with secondary antibodies (Alexa Fluor 488, Invitrogen A11055; Alexa Fluor 555, Invitrogen A31570; Alexa Fluor 647, Invitrogen A31573) diluted 1:500 in PBS containing 0.1% Tween-20 at RT for 1 h. The sections were washed three times for 5 min each at RT in PBS, followed by Hoechst 33258 (Invitrogen, H3569) staining for 10 min at RT. Ater three washes for 5 min each at RT in PBS, sections were mounted in EverBrite TrueBlack Hardset Mounting Medium (Biotium, 23018) and visualized using a Leica STELLARIS 5 microscope with a 40x objective.

#### Ambient RNA analysis

SoupX^21^ was used to estimate the levels of ambient RNA in all snRNA-seq dataset. Briefly, the automated method provided in SoupX was used for estimating the ambient RNA contaminated fraction and adjustCounts() was used to compute the final adjustment of RNA expression count matrix based on the estimated RNA contamination profile.

#### Western blotting analyses

Protein lysates were prepared from 10 mg frozen frontal cortical tissues using RIPA lysis buffer (50 mM Tris-HCl pH 8.0, 150 mM NaCl, 1 mM EDTA pH 8.0, 0.1% SDS, 0.5% Na-Deoxycholate, 1% Triton X-100) with protease inhibitor (Roche, 6127000) and homogenized using a Dounce homogenizer, followed by sonication (Diagenode Bioruptur 300, 5 cycles of 20 second on/off with MAX power). Protein concentrations were determined using a BCA protein assay (Thermo Fisher Scientific, 23227). Equal amounts of 10 μg protein lysate were loaded and separated on a 4-12% SurePAGETM gel (GenScript) and transferred to a polyvinylidene difluoride membrane (Millipore). Membranes were blocked in 5% non-fat dry milk buffer made in PBS-T (137 mM NaCl, 2.7 mM KCl, 10 mM Na2HPO4, 2 mM KH2PO4, adjusted to pH 7.4, 0.1%Tween 20) and probed with primary antibody over-night at 4 °C. Primary antibodies against total S6 ribosomal protein (Cell Signaling, 2217, 1:1000), phosphorylated S6 ribosomal protein (ser235/236) (Cell Signaling, 4858, 1:1000), and GAPDH (Cell Signaling, 2118, 1:3000) were used. After extensive washing with PBS-T, HRP-conjugated secondary antibodies (Abcam, ab6721) were added and incubated for 1 h at room temperature. SuperSignal West Femto Maximum Sensitivity Substrate (Thermo Scientific, 34094) was used to develop the signal following the manufacture’s protocol. Images were acquired with a Biorad ChemiDoc Touch Imaging System (Biorad) using the extended dynamic range to acquire images without saturation. Quantification was performed using ImageJ (version 1.54g).

#### Weighted correlation network analysis on neuronal cell clusters

WGCNA^99^ (weighted gene co-expression network analysis) was used to identify gene coexpression networks of neuronal clusters in the Emory cohort. This method identifies highly correlated gene clusters (termed modules) via unsupervised clustering. Pseudo-bulk expression for each excitatory and inhibitory neuronal cell cluster (n=10 and n=6, respectively) was analyzed separately with WGCNA using default parameters. Only differentially expressed genes in neuronal clusters were used for WGCNA analysis and a soft threshold power of 9 was used when constructing the network using blockwiseModules(). Hub genes were identified using signedKME() for each module. Correlation analysis between WGCNA modules and disease progression by grouping levels of pTDP-43 as described above was done using linear models on each module with Limma^100^ with multiple testing corrections, and the correlation was considered significant if p-adj ≤0.01. Gene ontology enrichment for each module was performed using clusterProfiler^101^ using the protein-coding hub genes with kME value > 0.8.

#### Analysis of cell-cycle scoring

Cell cycle analysis was performed following the default vignette in Seurat^18^ with the list of cell cycle markers^102^. The gene expression matrix was extracted from ArchR’s ArchRProject to create the Seurat object before using Seurat’s CellCycleScoring().

#### Differential cell abundance analysis

To identify differences in cell composition across the donor groups with different levels of pTDP-43 in each cell cluster in each major cell type, we calculated the relative percentage of each cluster in each major cell type for each sample. The differential cell proportions were estimated using Kruskal-Wallis test with Benjamini-Hochberg correction comparing the control with different pTDP-43 level groups. P-values > 0.05 were considered not significant. For the neuronal clusters, the significance of differential abundance was further analyzed using MASC^103^, which considers the mixed-effect model with a binomial distribution accounting for technical confounders and biological variation. We included the following fixed covariates in the model: sex, sample status (control and *C9orf72* ALS/FTD cases), and level of pTDP-43. Cell clusters were considered significant at FDR-adjusted P < 0.05 and absolute odds ratio >0. The results of MASC analysis are shown in **Fig. 6g**.

#### Reverse deconvolution with pTDP-43 sorted bulk RNA-seq datasets

Published RNA-seq datasets of FACS sorted pTDP-43 NeuN+ neurons from the frontal cortex of *C9orf72* ALS/FTD donors were downloaded from the Gene Expression Omnibus (GEO) database under accession GSE126543^77^. Pseudo-bulk snRNA-seq data for each cell cluster were analyzed using CIBERSORTx^78^ with default parameters.

## Acknowledgements

We would like to thank Dr. Cynthia Vied at the Translational Science Laboratory of Florida State University for help with Illumina sequencing; Dr. Steven Sloan from the Department of Human Genetics at Emory University for assistance with the initial stage of astrocyte lineage analysis; Dr. Ryan Corces at the Gladstone Institute of Neurological Disease for assistance in the initial analysis of snATAC-seq data. This work was supported by U.S. Public Health Service Awards R35 GM139408 (VGC); R35 NS111602, R01 HG008935, U01 MH116441 (PJ) from the National Institutes of Health. H-LW was supported by NIH F32 ES031827. The content is solely the responsibility of the authors and does not necessarily represent the official views of the National Institutes of Health.

## Author contributions

H-LW, VGC and ZTM conceived, designed the project, and wrote the manuscript. H-LW planned and performed multiome experiments and analyzed data; AMV and TFG performed experiments to quantify pTDP-43 levels; MG performed analyses of cortical tissue pathology for the Emory samples; JDG recruited donors and obtained clinical information for the Emory samples; MEM and MGT performed neuropathological assessments and tissue dissections for the Mayo samples; PJ planned experiments; CY and JH analyzed the MERSCOPE images; JFZ performed Western blot analyses.

## Data availability

All data generated in this work are available through GEO accession number GSE212630. Reviewers can access these data using token yruhcwmcrfodrkn. All scripts used for analyzing the data in this manuscript can be found on the GitHub repository, https://github.com/wanghlv/c9alsftd_multiome.

