## Supplementary information for "pTDP-43 levels correlate with cell type specific molecular alterations in the prefrontal cortex of *C9orf72* ALS/FTD patients"

**Supplementary Figures**

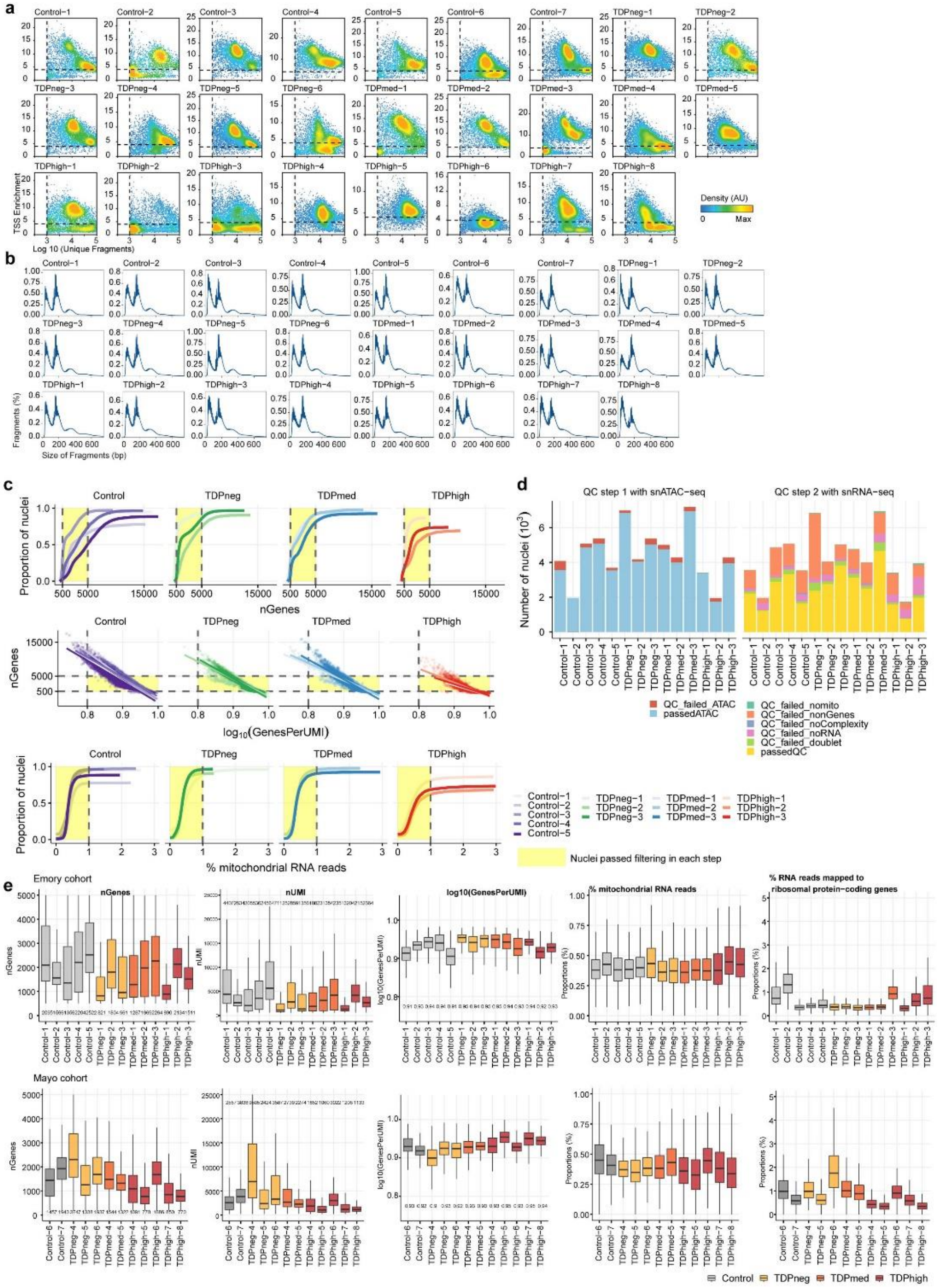


**Supplementary Fig. 1 Quality control metrics of the paired multiomics snATAC-seq and snRNA-seq data from the same nuclei of all donor samples in both Emory and Mayo cohorts.** Quality control metrics for the snATAC-seq dataset showing (a) the correlations between TSS enrichment score and unique nuclear fragments per cell, and (b) the distribution of fragment size for cells in each sample passing ArchR QC thresholds. Nuclei passing snATAC-seq quality control (TSS >4, number of fragments >1000, and doublet removal) are depicted in the upper quadrant in (a). (c) Cumulative proportion of detected genes (nGenes), log10 transformed genes per UMI, and the percentage of reads mapped to the mitochondrial genome are presented for each snRNA-seq sample within the Emory cohort, categorized by pTDP-43 levels. The yellow shaded area denotes nuclei that passed QC filtering based on snRNA-seq data (nGenes: 500-5000; log10(GenesPerUMI) > 0.8, % reads mapped to the mitochondria genome is < 1%). (d) Quantification of the removal of low-quality nuclei at each quality control step using paired snATAC-seq and snRNA-seq data for every sample in the Emory cohort, relative to the yellow-marked nuclei that passed QC. (e) Quality control matrices for the snRNA-seq datasets for nuclei that passed filtering for samples in the Emory (top) and Mayo (bottom) cohorts.

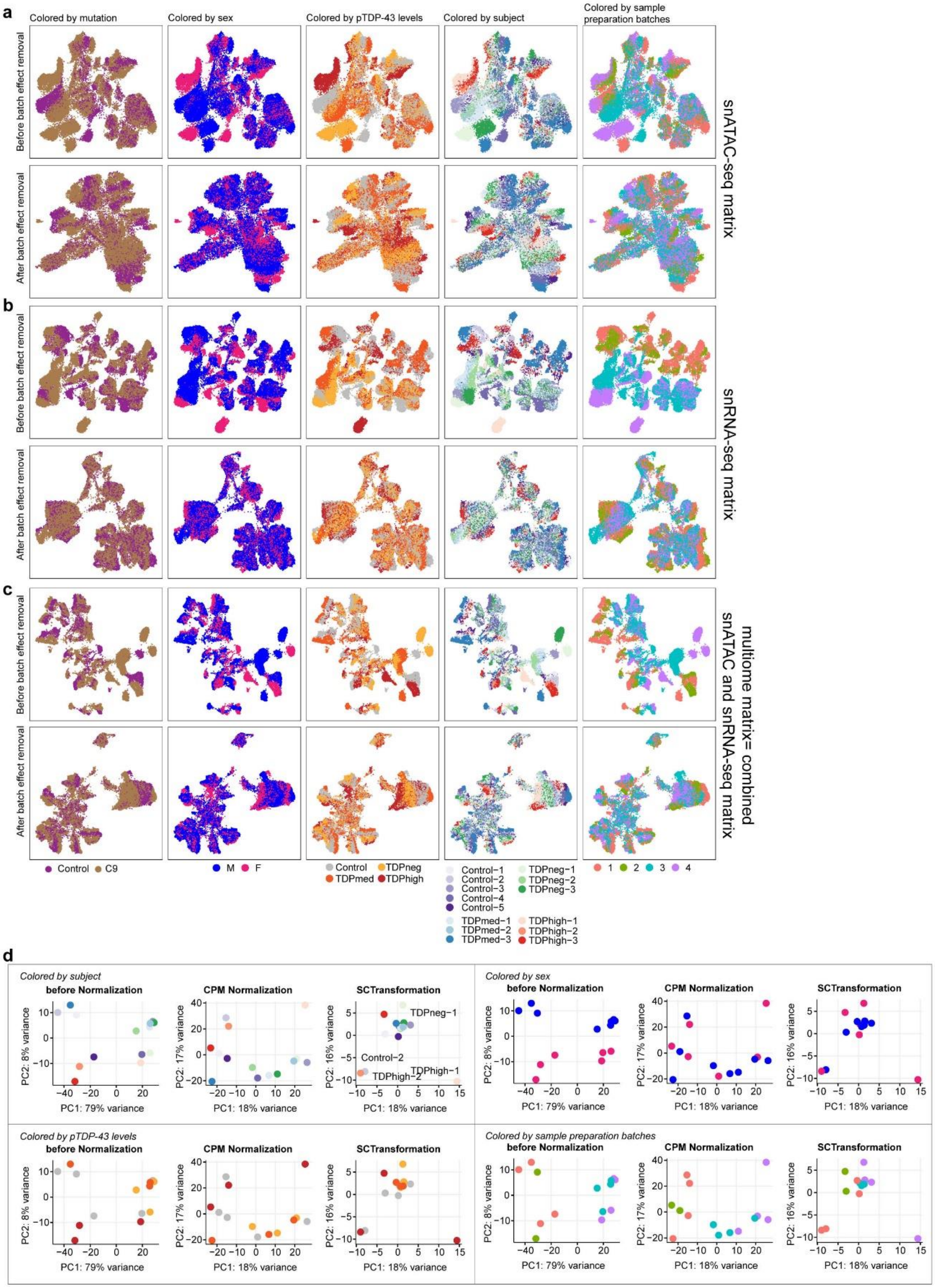


**Supplementary Fig. 2. UMAP visualizations before and after batch effect correction for samples in the Emory cohort.**

(a) UMAP plots, colored by mutation, sex, pTDP-43 levels, subjects, and sample preparation batches were generated from snATAC-seq. (b) UMAP plots, colored by mutation, sex, pTDP-43 levels, subjects, and sample preparation batches were generated from snRNA-seq. (c) UMAP plots, colored by mutation, sex, pTDP-43 levels, subjects, and sample preparation batches were generated from combined multiomics datasets. The same color legend is shared between panels a, b, and c. (d) PCA plots of snRNA-seq datasets before and after normalization.

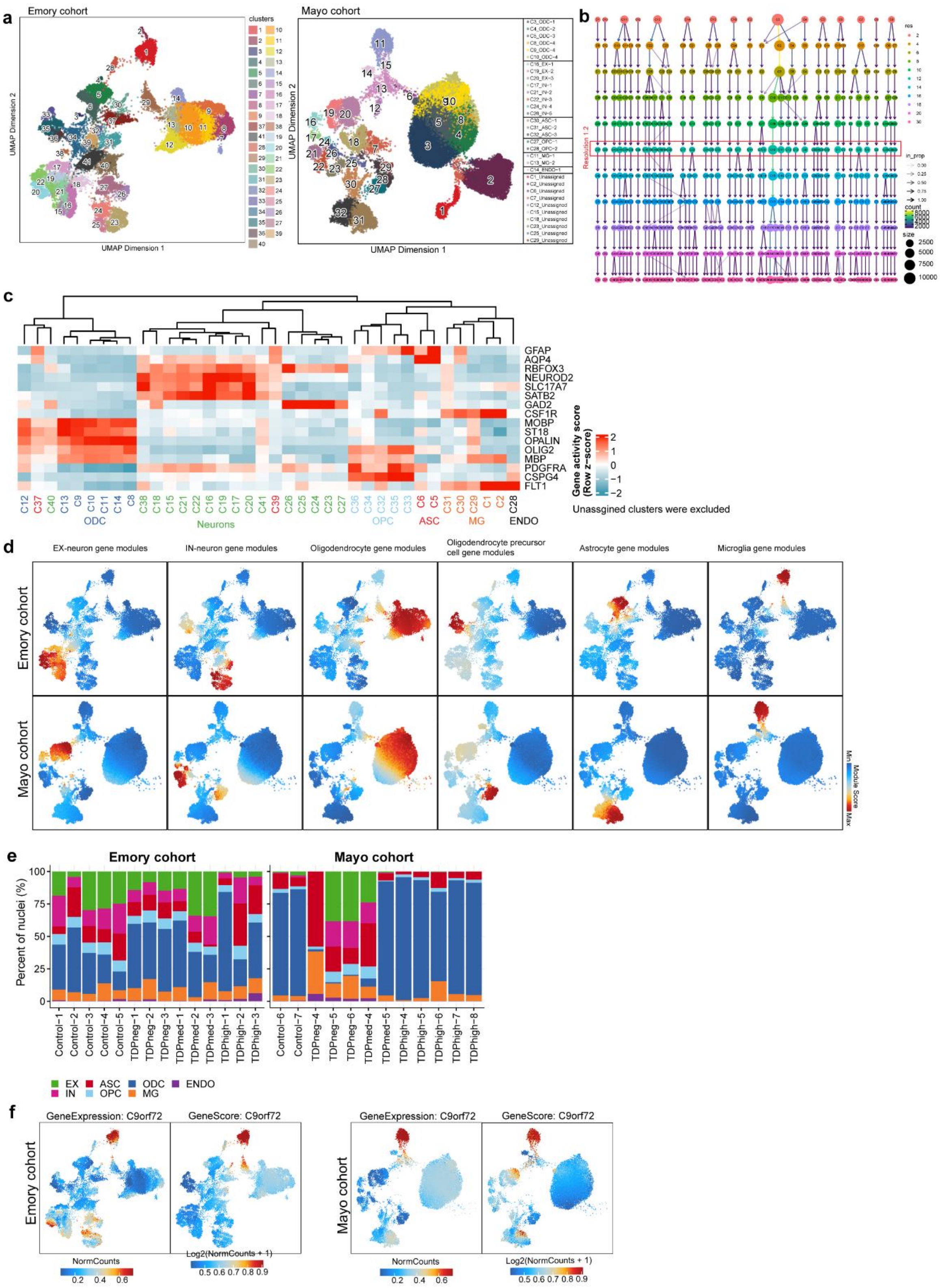


**Supplementary Fig. 3. Quality control, gene module and gene activity scores.** (a) UMAP plot colored by clusters for Emory and Mayo cohorts. (b) Cluster stability was assessed using Clustree analysis with increasing resolutions from 0.2 to 2 in the Emory cohort dataset. Resolution 1.2 was selected and highlighted. (c) Row-normalized single-nucleus gene score heatmap (left) and violin plots (right) of cell-type marker genes in each cluster in the Emory cohort. (d) Gene module scores of major cell types, excluding endothelial cells, for the Emory (top row) and Mayo (bottom row) cohorts. (e) Proportions of major cell types for *C9orf72* ALS/FTD cases staged by pTDP-43 abundance. (f) Gene activity score and gene expression of the *C9orf72* gene for the Emory (left) and Mayo (right) cohorts.

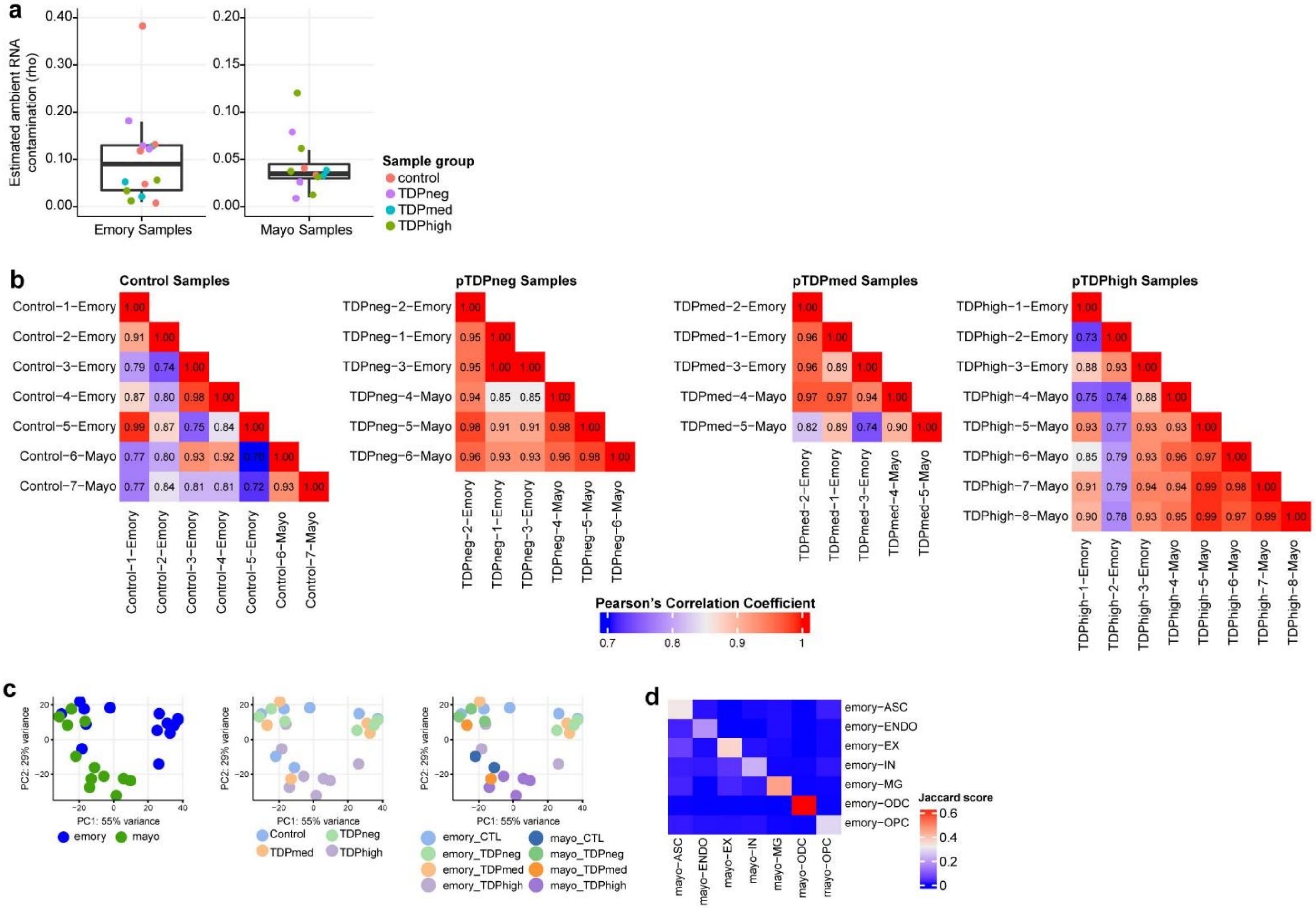


**Supplementary Fig. 4. Ambient RNA analysis and sample correlations between two brain banks.** (a) Average ambient RNA contamination in each snRNA-seq library. All libraries have less than 20% estimated global ambient RNA contamination except for control library Control-4 from the Emory cohort. (b) Pearson correlation analysis of samples in control, pTDPneg, pTDPmed, and pTDPhigh groups from both brain banks. (c) PCA plots of snRNA-seq datasets from Emory and Mayo samples. (d) Heatmap showing the similarities of marker genes between cortical cell types from Emory and Mayo samples. The jaccard score indicates the percentage of pairwise overlapping genes, and the ODC shows the greatest similarity between the two cohorts.

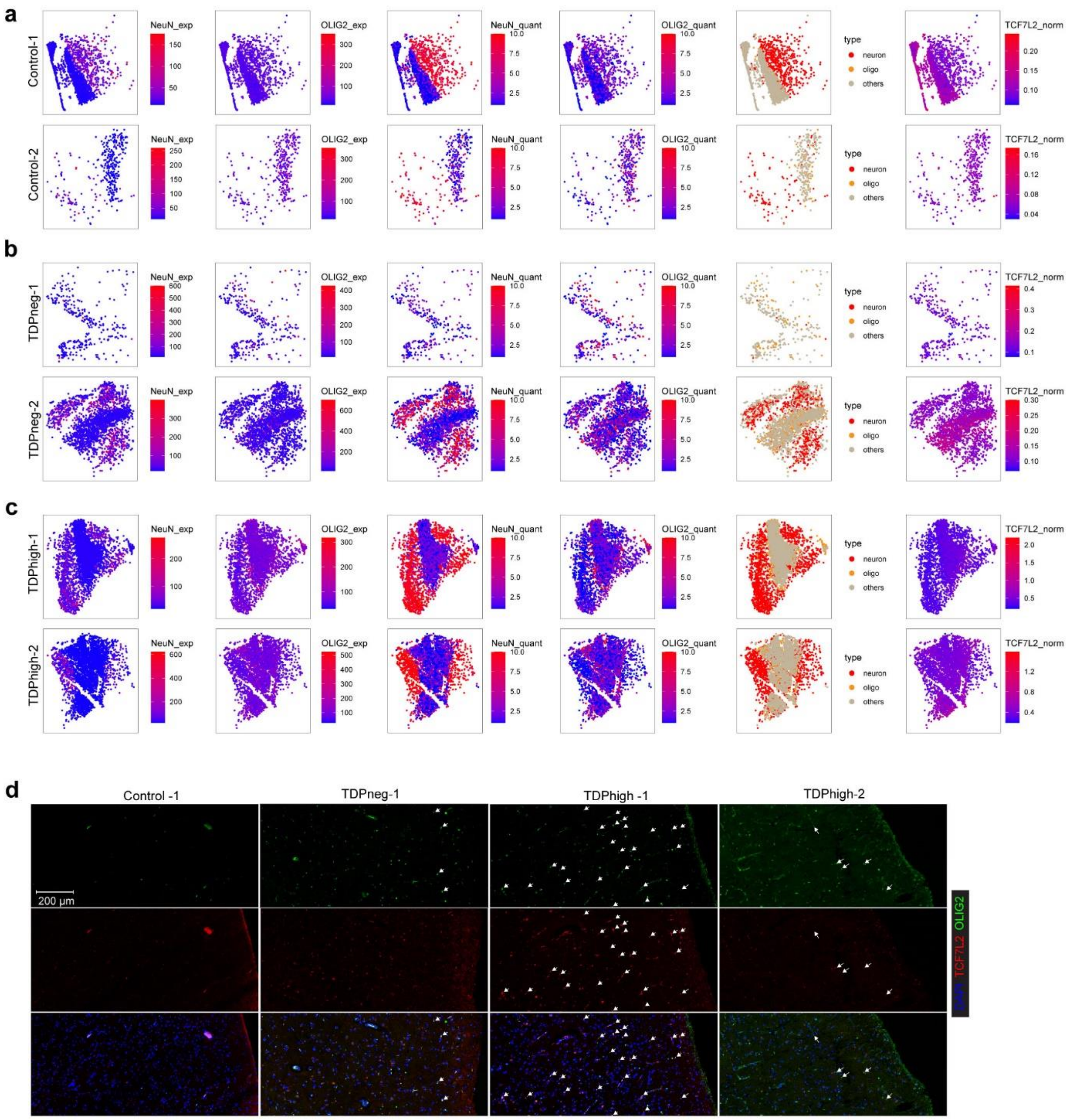


**Supplementary Fig. 5. OLIG2+/TCF7L2+ nuclei are present in high abundance in TDPhigh samples.** (a-c) The mean signal intensities for each segmented nucleus were shown for immunostaining that was done using the MERSCOPE. Results from two control samples are shown in panel (a); results from two TDPneg samples are shown in panel (b); and results from two TDPhigh samples are shown in panel (c). For each sample from left to right the mean signal intensity is shown for NeuN and OLIG2 staining as NeuN_exp and OLIG2_exp, respectively. The quantile normalized value for NeuN and OLIG2 staining are shown as NeuN_quant and OLIG2_quant, respectively. The classification of nuclei as neurons, oligodendrocytes (oligo), and others are also shown for each sample. The last panels on the far right show the normalized TCF7L2 intensity by DAPI intensity for samples from control, pTDPneg and pTDPhigh groups. (d) Images from immunofluorescence microscopy of TCF7L2 and OLIG2 using 40 µm sections followed by confocal microscopy.

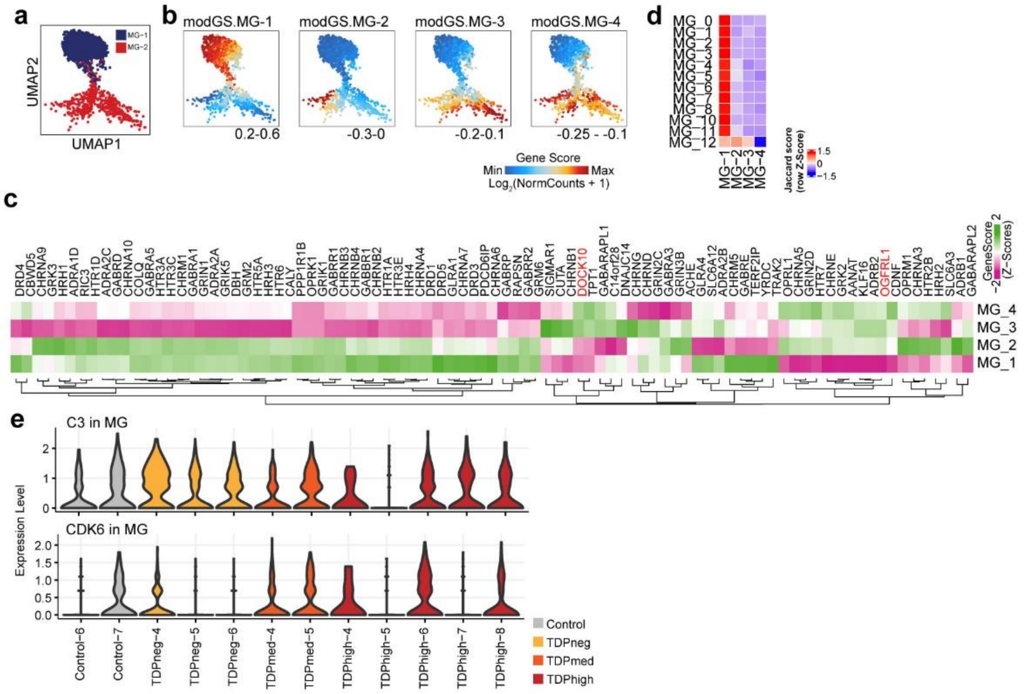


**Supplementary Fig. 6. Additional data from analysis of microglia.**
(a) UMAP plots of microglia clusters in Mayo samples. (b) Same UMAP as in (a) colored by expression of cluster specific genes, rendered with imputation of ATAC-seq module gene scores; module genes are listed in Fig. 4b. (c) Gene score activities of neurotransmitter receptors in each microglia cluster. (d) Heatmap showing the similarities of marker genes between Sun et al. and all microglia clusters found in this study. (e) Violin plots showing gene expression levels of the *C3* and *CDK6* genes in microglia of Mayo samples.

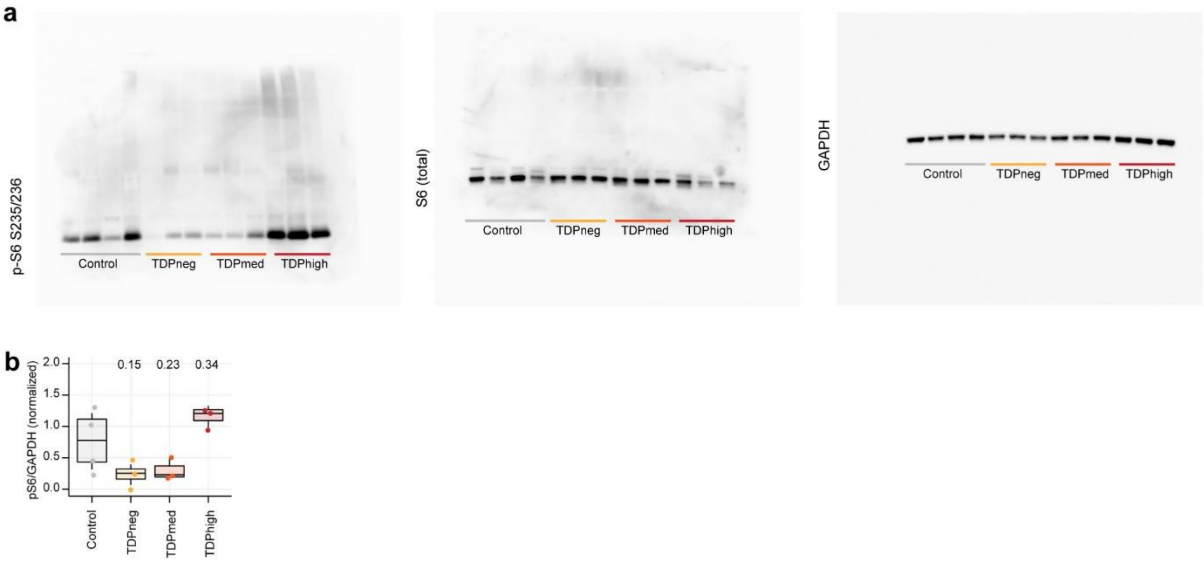


**Supplementary Fig. 7. Analysis of changes in phosphorylation of ribosomal protein S6.** (a) Full size images of western blots used to obtain the cropped images shown in Fig. 5h. (b) Quantification of the ratio of phosphorylated ribosomal protein S6 to GAPDH. 1-way ANOVA with Tukey's post hoc test, adjusted p-values are shown.

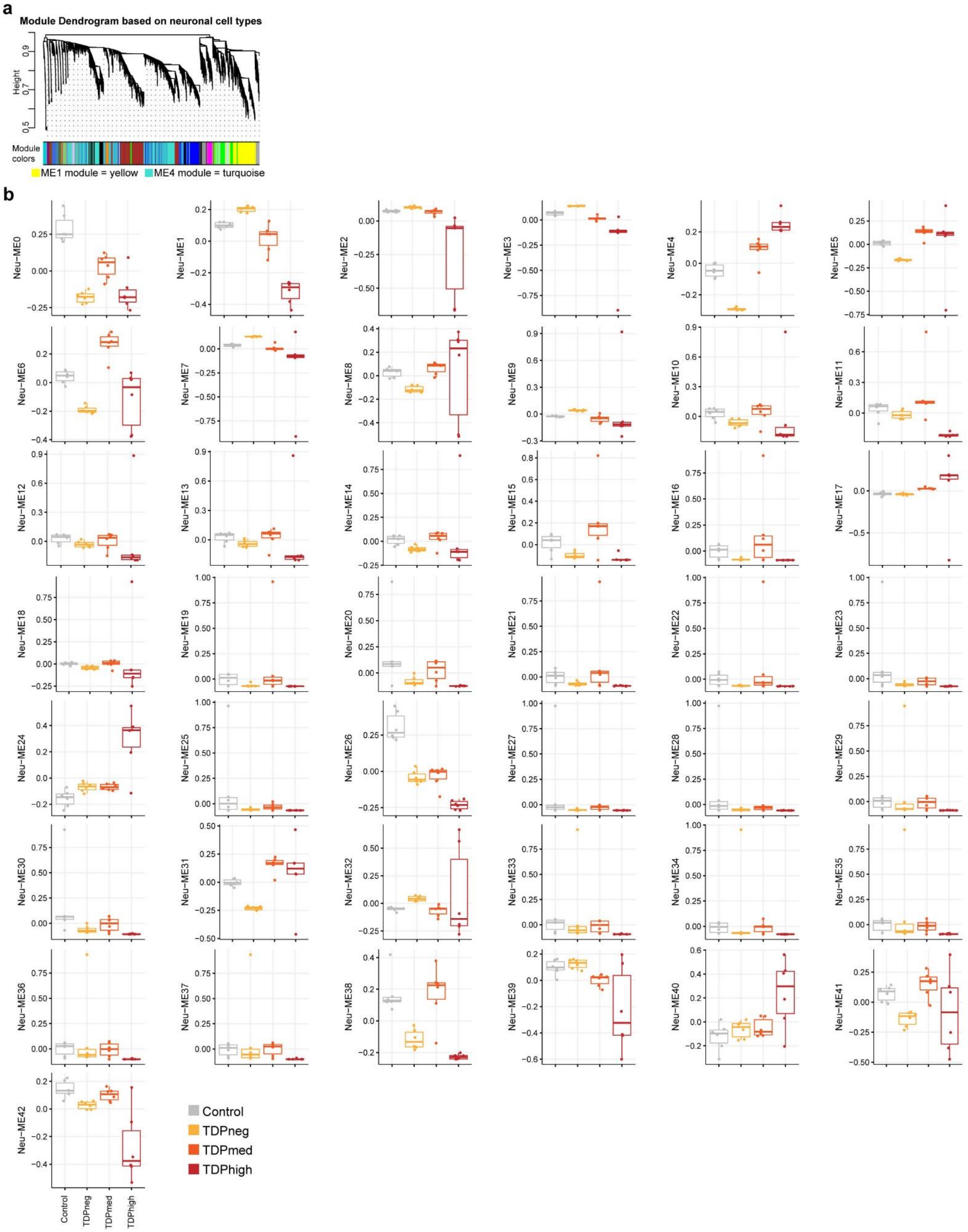


**Supplementary Fig. 8. WGCNA analysis of neuronal clusters**

(a) WGCNA module dendrograms. (b) Significance of WGCNA modules with different levels of pTDP-43.

**Supplementary Tables**

**Table 1**. Sample metadata.
**Table 2**. Single cell multiome snATAC+snRNA quality control data.
**Table 3**. List of marker genes for each cell cluster.
**Table 4**. List of differentially expressed genes and differential chromatin accessible regions.
**Table 6**. List of WGCNA module eigengenes found in neuronal clusters.
